# A silent disco: Differential effects of beat-based and pattern-based temporal expectations on persistent entrainment of low-frequency neural oscillations

**DOI:** 10.1101/2020.01.08.899278

**Authors:** Fleur L. Bouwer, Johannes J. Fahrenfort, Samantha K. Millard, Niels A. Kloosterman, Heleen A. Slagter

## Abstract

The brain uses temporal structure in the environment, like rhythm in music and speech, to predict the timing of events, thereby optimizing their processing and perception. Temporal expectations can be grounded in different aspects of the input structure, such as a regular beat or a predictable pattern. One influential account posits that a generic mechanism underlies beat-based and pattern-based expectations, namely entrainment of low frequency neural oscillations to rhythmic input, while other accounts assume different underlying neural mechanisms. Here, we addressed this outstanding issue by examining EEG activity and behavioral responses during silent periods following rhythmic auditory sequences. We measured responses outlasting the rhythms both to avoid confounding the EEG analyses with evoked responses, and to directly test whether beat-based and pattern-based expectations persist beyond stimulation, as predicted by entrainment theories. To properly disentangle beat-based and pattern-based expectations, which often occur simultaneously, we used non-isochronous rhythms with a beat, a predictable pattern, or random timing. In Experiment 1 (N = 32), beat-based expectations affected behavioral ratings of probe events for two beat-cycles after the end of the rhythm. The effects of pattern-based expectations reflected expectations for one interval. In Experiment 2 (N = 27), using EEG, we found enhanced spectral power at the beat frequency for beat-based sequences both during listening and silence. For pattern-based sequences, enhanced power at a pattern-specific frequency was present during listening, but not silence. Moreover, we found a difference in the evoked signal following pattern-based and beat-based sequences. Finally, we show how multivariate pattern decoding and multi scale entropy – measures sensitive to non-oscillatory components of the signal – can be used to probe temporal expectations. Together, our results suggest that the input structure used to form temporal expectations may affect the associated neural mechanisms. We suggest climbing activity and low-frequency oscillations may be differentially associated with pattern-based and beat-based expectations.

## Introduction

Predicting the timing of incoming events optimizes processing in our dynamic environment (Nobre & van Ede, 2018), as it allows the brain to increase sensitivity to events at predicted times (Auksztulewicz et al., 2019), without the need for constant vigilance (Breska & Deouell, 2017; Rimmele et al., 2018; Schroeder & Lakatos, 2009a, 2009b). Entrainment models (Large & Jones, 1999) provide a mechanistic explanation for temporal expectations, by assuming that the phase and period of low-frequency neural oscillations synchronize to external rhythmic stimulation, causing optimal neural excitability at expected times (Haegens & Zion Golumbic, 2018; Henry & Herrmann, 2014; Schroeder & Lakatos, 2009b). In line with this, behavioral performance is improved for events in phase with an external rhythm (Bouwer & Honing, 2015; Herbst et al., 2022; Jones et al., 2002; Large & Jones, 1999), behavioral responses depend on the phase of delta oscillations (Arnal et al., 2014; Cravo et al., 2013; Henry et al., 2014; Henry & Obleser, 2012), and low frequency oscillations phase lock to rhythmic input (Doelling et al., 2019; Nozaradan et al., 2011; Stefanics et al., 2010). Note that in the current study, we focus on such perceptual entrainment – the putative alignment of low-frequency oscillations to an external stimulus and how this is associated with temporal expectations – and not on the broader connotation of entrainment that includes motor synchronization to rhythm (Damm et al., 2019; Repp, 2005).

Entrainment has mainly been studied in the context of periodic (“beat-based”) sensory input, but temporal expectations can also be based on memory for absolute intervals (Breska & Deouell, 2017; Breska & Ivry, 2016; Morillon et al., 2016; Teki et al., 2011), either in isolation (“cue-based”), or as part of a predictable pattern of intervals (“pattern-based”, see (Nobre & van Ede, 2018)). Predictable temporal patterns may be especially important in speech and non-Western music, which is not necessarily based on a metrical structure with a prominent periodicity in the range of human beat-based perception (London, 2012; London et al., 2017; Polak et al., 2016). Expectations based on predictable patterns in aperiodic sequences afford similar behavioral benefits as expectations based on a beat (Bouwer et al., 2020; Heideman et al., 2018; O’Reilly et al., 2008), but pose a possible challenge for entrainment models, which are arguably better suited to explain temporal expectations for periodic input (Breska & Deouell, 2017; Rimmele et al., 2018). Some have suggested that entrainment models can account for pattern-based expectations by assuming multiple coupled oscillators at different frequencies and with different phases (Tichko & Large, 2019), or by assuming flexible top-down phase resets at expected moments, though this would entail some top-down mechanism, making observed entrainment the consequence, rather than the cause of expectations (Meyer et al., 2019; Obleser & Kayser, 2019; Rimmele et al., 2018).

Alternatively, however, pattern-based and beat-based expectations could arise from dissociable neural mechanisms. For cue-based expectations, tentative evidence for a different underlying mechanism comes from a series of studies looking at the contingent negative variation (CNV), an event-related potential (ERP) component that peaks at expected moments (Praamstra et al., 2006). The CNV resolved faster for beat-based than cue-based expectations (Breska & Deouell, 2017), and cerebellar patients showed selective impairments in forming cue-based, but not beat-based expectations (Breska & Ivry, 2018, 2020). However, in these studies, the intended beat-based sequences were isochronous. Isochronous sequences can, in addition to a beat, elicit temporal expectations through learning the repeated, identical interval (Breska & Ivry, 2016; Keele et al., 1989). Thus, the differences in responses may here be explained by more precise cue-based or pattern-based expectations in the isochronous, beat-based condition. Moreover, these studies tested temporal expectations based on the contingency between a cue and an interval (e.g., learning a single interval), and it is unclear whether temporal expectations based on patterns are based on the same mechanism (Nobre & van Ede, 2018). In our own recent work, we specifically compared beat-based and pattern-based expectations, and we found no difference in the effects of these expectations on early auditory ERP responses, suggestive of similar modulation of sensory processing (Bouwer et al., 2020), but we observed suppression of sensory processing of unexpected events in beat-based rhythms, even when these events were fully predictable based on their pattern, suggesting different underlying mechanisms (Bouwer et al., 2020). However, in this study, we did not directly probe the neural mechanisms underlying beat-based and pattern-based expectations, rendering it unclear whether they are subserved by shared or separate neural dynamics.

In the current study, we directly examined the role of entrainment in beat-based and pattern-based expectations, using non-isochronous rhythms designed to properly disentangle these two types of expectations. When studying entrainment, an important challenge has been to differentiate between real entrainment (“in the narrow sense”, see (Obleser & Kayser, 2019) and regular evoked potentials, or similar phase locked responses that resemble entrainment with common analysis techniques (Zoefel et al., 2018), and that may not differentiate between beat-based and memory-based expectations (Breska & Deouell, 2017). Crucially, to sidestep these issues, here we examined responses in a silent window after cessation of the rhythmic input, directly testing the prediction of entrainment models that entrainment should outlast sensory stimulation (Haegens & Zion Golumbic, 2018; Obleser & Kayser, 2019; Pesnot Lerousseau et al., 2021; Zoefel et al., 2018).

Behaviorally, the effects of persistent entrainment on perception have been shown for auditory rhythm (Hickok et al., 2015; Jones et al., 2002), though this effect is not always found (Bauer et al., 2015; Lin et al., 2021), possibly due to heterogeneity in the population, effects of musical training, and the intricacies of stimulus design (Assaneo et al., 2019; Cameron & Grahn, 2014; Saberi & Hickok, 2022a; Sun et al., 2021). At a neural level, several studies reported persistent entrainment in the visual (de Graaf et al., 2013; Mathewson et al., 2012), and auditory (Kösem et al., 2018; Pesnot Lerousseau et al., 2021; van Bree et al., 2021; Wilsch et al., 2020) domain (see (Saberi & Hickok, 2022b) for an overview). However, in these studies, isochronous stimuli were used, making it unclear whether the expectations probed were based on a beat, or were formed based on the single repeating interval (i.e., cue-or pattern-based). Moreover, persistent entrainment was not specific to the frequency of the input (Wilsch et al., 2020), or only occurred in the gamma (Pesnot Lerousseau et al., 2021), or alpha ranges (de Graaf et al., 2013; Mathewson et al., 2012), while humans have a preference for forming temporal expectations at slower rates (Ding et al., 2017; Merchant et al., 2015; Zalta et al., 2020), as naturally present in speech and music (i.e., the delta and theta range). Thus, not only is evidence for whether entrainment can account for pattern-based expectations lacking, evidence for persistent neural entrainment in response to beat-based rhythms remains elusive as well.

In the current study, participants listened to non-isochronous auditory sequences (similar to those used in (Bouwer et al., 2020)) with either a regular beat (eliciting beat-based expectations), a predictable pattern (eliciting pattern-based expectations), or random timing (no expectations). The non-isochronous beat-based sequences had a varying surface structure, similar to patterns used to probe beat-based processing in many neuroimaging (Grahn & Brett, 2007; Grahn & Rowe, 2009; Leow & Grahn, 2014), behavioral (Bouwer et al., 2018, 2021; Cameron & Grahn, 2014; Povel & Essens, 1985), and electrophysiological studies (Lenc et al., 2021). Therefore, the beat could not be extracted from the rhythmic signal by simply learning the transition of temporal intervals – as is possible in isochronous sequences.

Each sequence was followed by a silent period. In Experiment 1, we asked participants to rate how well probe tones, presented at various time points during the silent period, fitted the preceding sequence. We expected the ratings to be affected by both the beat-based and pattern-based expectations elicited by the sequences. In Experiment 2, we recorded EEG activity both during presentation of the sequences and during the silence. If entrainment underlies temporal expectations, we should see persistent power at the frequency of the beat or pattern during the silence. Alternatively, if climbing activity in the form of a CNV underlies temporal expectations, we should see a CNV peaking at expected time points in the silence. As a CNV is generally contingent on a preceding stimulus (Kononowicz & Penney, 2016), we expect the CNV to only occur once after cessation of the rhythm, with its timing dependent on the one interval that is expected at that point, and therefore, to be dissociable from persistent power at frequencies prominent in the overall rhythmic pattern.

In addition to examining spectral power and evoked responses, we explored indexing temporal expectations in the silence using multi scale entropy (MSE) – a measure of signal irregularity (Kosciessa et al., 2020) – and multivariate pattern decoding. While MSE provides us with a method to look at possible non-sinusoidal contributions to the EEG signal, decoding allows us to look at how entrainment evolves over time. As recently argued, the oscillatory dynamics underlying temporal expectations may be subject to changes in power and frequency, depending on coupling between sound and brain, and on the properties of the neural dynamics themselves (Doelling & Assaneo, 2021). Once the rhythmic sensory input ceases, the oscillatory activity in the brain may quickly return to an intrinsic resonance frequency (Doelling & Assaneo, 2021). MSE and decoding may provide useful tools to study these neural dynamics. Note that we consider these analyses exploratory in nature, and results should be interpreted as such.

## Materials and methods

### Participants

Thirty-two participants (18 women), aged between 18 and 44 years old (M = 24, SD = 5.6) took part in the behavioral Experiment 1, and 32 participants (26 women), aged between 19 and 28 years old (M = 23, SD = 2.5) took part in the EEG experiment (Experiment 2), in exchange for course credit or monetary compensation. Due to technical problems, the EEG data from five participants was not recorded correctly, hence we report the results for 27 participants (21 women, between 19 and 28 years old, M = 23, SD = 2.4). For the behavioral experiments, we used mixed-effects models, which need both the number of participants and the number of items to be taken into account to assess power (Brysbaert & Stevens, 2018). In two similar experiments in which ratings in response to rhythms of varying complexity were analyzed, small-sized effects were replicated with a total number of around 275 responses per condition (Bouwer et al., 2018). For our new experimental paradigm, we included a multiple of this number of trials (960 responses per condition and probe position in Experiment 1 – 32 participants with 30 responses each in each cell – and 486 responses per condition and probe position in Experiment 2 – 27 participants with 18 responses). Previous EEG experiments examining persistent entrainment in the auditory domain used sample sizes ranging from fifteen (Pesnot Lerousseau et al., 2021) to twenty-one (van Bree et al., 2021), similar to the sample size used in a study looking at different types of temporal expectations (Breska & Deouell, 2017). Here, we report data from twenty-seven participants, thus exceeded typical sample sizes used previously. None of the participants reported a history of hearing or neurological problems, and all provided written informed consent prior to the onset of the study. The experiment was approved by the Ethics Review Board of the Faculty of Social and Behavioral Sciences of the University of Amsterdam.

### Stimuli

We used patterns marked by woodblock sounds of 60 ms length, generated in GarageBand (Apple Inc.), as previously used in (Bouwer et al., 2020) to elicit beat-based and pattern-based expectations (Figure 1). Each pattern was 1800 ms long and consisted of five temporal intervals. The number of tones was chosen to be within the range that was previously shown to allow for learning of a predictable pattern (Schultz et al., 2013), while the length of the pattern was such that the formation of beat-based expectations with a period of the entire pattern would be unlikely in the pattern-based sequences, given that this would require hearing a beat at 0.55 Hz, which is very far from the range at which humans can typically perceive a beat (Honing & Bouwer, 2019; London, 2012). Sequences (beat-based, pattern-based, or with random timing) were constructed by concatenating four patterns and a final tone, for a total sequence length of 7260 ms (four patterns of 1800 ms, plus 60 ms for the final tone).

**Figure 1.**
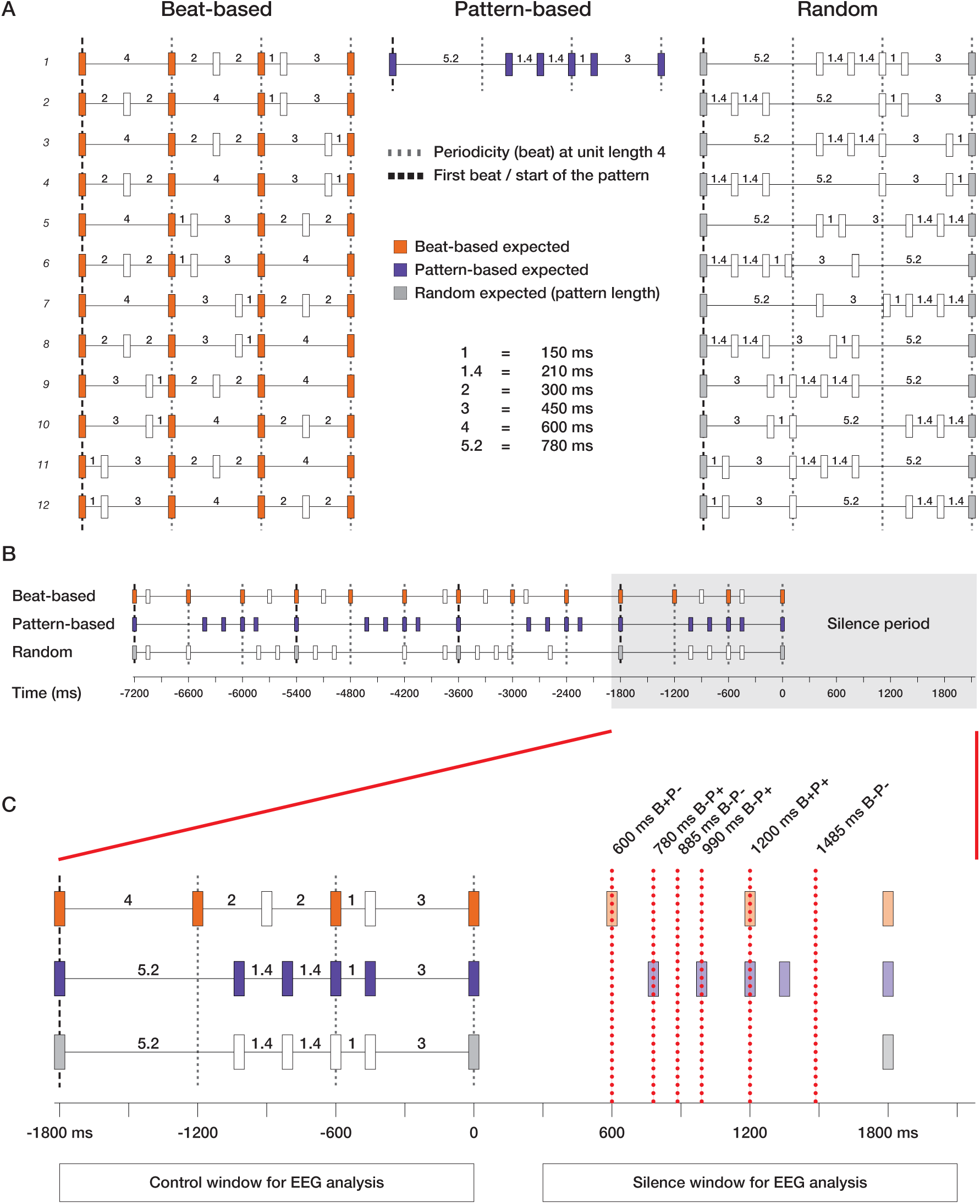
Schematic overview of the rhythmic stimuli used and the task. A) Twelve patterns of five temporal intervals with integer ratio durations and an event at each 600 ms period were created to form beat-based sequences. Equivalent patterns without a regular beat every 600 ms were created by using non-integer ratio durations, while keeping the number of intervals and grouping structure the same (random condition). For the pattern-based sequences, only pattern 1 was used, to allow for learning of the intervals. B) Four semi-randomly chosen patterns were concatenated to form rhythmic sequences. In both the beat-based and random sequences, the last pattern was always pattern 1 or 2, to equate the acoustic context preceding the silent period. C) To measure behavioral effects of expectations, a probe tone could appear at various temporal positions in the silent period (indicated by the dashed red lines), predictable based on a beat (B+, light orange), predictable based on the pattern (P+, light purple), or unpredictable based on the beat (B-) or pattern (P-). Subjects had to indicate how well the probe tone fitted the preceding rhythm. In Experiment 1, all 6 probe tone positions were used. In Experiment 2, only the last 3 probe positions were used.

In the twelve patterns used to create beat-based sequences (Figure 1), temporal intervals were related to each other with integer-ratio durations. The shortest interval had a length of 150 ms, with the relation between the five intervals used of 1:2:2:3:4 (i.e., 150, 300, 300, 450, and 600 ms). The sounds were grouped such that a perceptually accented sound (Povel & Okkerman, 1981) occurred every 600 ms (every unit length 4), giving rise to a beat at 100 beats per minute, or 1.67 Hz, within the range of preferred tempo for humans (London, 2012). All beat-based patterns were strictly metric, with the beat always marked by a sound (Grahn & Brett, 2007). Sequences of beat-based patterns were constructed from four semi-randomly chosen patterns, with the restriction that the last pattern of the sequences was always pattern 1 or 2 (see Figure 1). This way, the final 600 ms preceding the silence epoch was equated in terms of the acoustic context, to make the bleed of auditory ERPs into the silence as similar between conditions as possible. Note that in beat-based sequences, a sound could be expected every 600 ms based on the beat, but the surface structure of the pattern was unpredictable, due to the random concatenation of patterns.

To create patterns that did not allow for beat-based expectations (“aperiodic” patterns, see Figure 1), the ratios by which the temporal intervals were related were changed to be non-integer (1:1.4:1.4:3:5.2, or 150, 210, 450, and 780 ms respectively). In these patterns, no marked beat was present at unit length four, nor at any other subdivision of the 1800 ms pattern (Bouwer et al., 2020), while the patterns were matched to their periodic counterparts in terms of overall length, event density, number of sounds, and grouping structure.

From the aperiodic patterns, two types of sequences were created: pattern-based and random sequences. To create sequences allowing for pattern-based expectations, we concatenated four identical patterns. To be able to use the data with an EEG-based decoding analysis (Experiment 2), we needed the timing of expectations in the silence to be identical for each sequence, hence we restricted the pattern-based sequences to only pattern 1. The use of a single pattern was not only necessary for decoding, but also optimized the experiment for pattern-based expectations, since participants only had to memorize one pattern, allowing them to easily form expectations, even if the single sequences were only four patterns long.

For the random sequences, four semi-randomly chosen aperiodic patterns were concatenated. Like for the beat-based sequences, the final pattern was always pattern 1 or 2, equating the final 600 ms of the sequences in terms of acoustics. In the random sequences, the timing of sounds could not be predicted based on the surface structure of the pattern, nor on the basis of an underlying beat.

A spectral analysis of the stimuli confirmed that in the beat-based sequences, a peak was present at the beat frequency of 1.67 Hz as well as at 3.33 Hz (see Figure 2). The 3.33 Hz peak is a harmonic of the beat frequency, but also the frequency at which participants may perceive subdivisions of the beat (e.g., an extra layer of perceived metrical regularity with a period of 300 ms). In the pattern-based and random sequences, peaks were more distributed, in line with the more irregular nature of these rhythms, and the highest peaks in the range in which a beat can normally be perceived (the delta-range, 0.5-4 Hz) were at 2.22 and 3.89 Hz.

**Figure 2.**
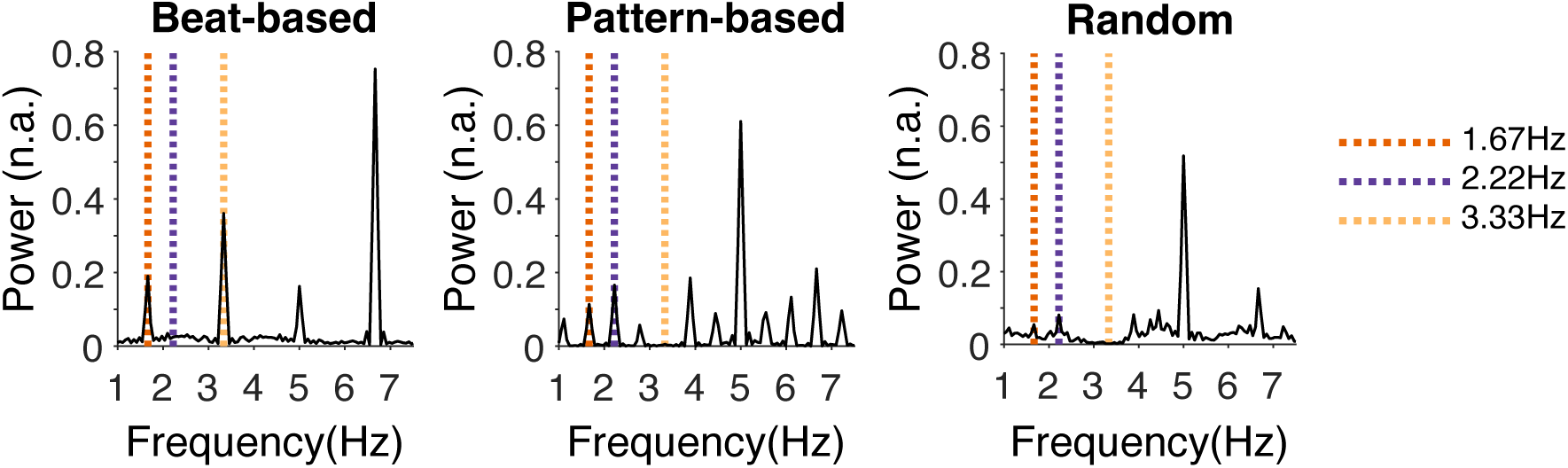
Spectral analysis of the sound signal of the different rhythmic sequences. Ten sequences per condition were generated to base the spectral analysis on. Note that these sequences were identical for the pattern-based condition, but semi-random for the beat-based and random conditions. The envelope of each sequence was obtained by performing a Hilbert transform, and subsequently, a fast fourier transform (fft) was used to obtain the spectral decomposition and power values were averaged over ten sequences. n.a.= normalized amplitude.

In Experiment 1, to assess the persistence of temporal expectations behaviorally, on each trial, a probe tone was presented at 600, 780, 885, 990, 1200, or 1485 ms after the onset of the last tone of the sequence (see Figure 1C), and participants provided ratings for how well probe tones fitted the preceding rhythm. These positions were carefully chosen to represent times at which a tone could be expected based on the beat (600, 1200 ms), based on memory for the pattern (780, 990, 1200 ms), or neither (885, 1485 ms). Note that the latter two probe tones that were unexpected based on the beat (780, 885, 990, and 1485 ms) did not fall on subdivisions of the beat.

Experiment 2 contained both trials in which a probe tone was presented (25% of the trials, using only the last three probe positions), and trials in which a 7260 ms sequence was followed by a silence period without a probe tone (75% of the trials). The latter were used for EEG analyses, uncontaminated by a probe presentation. The silent period lasted for 2100 ms after the onset of the last sound, providing 300 ms immediately following the final sound to allow for ERPs to mostly return to baseline, and 1800 ms (three full beat cycles, or a full cycle of the repeating pattern) of silence for the analysis. After the silence, the onset of the next sequence was jittered between 25 and 75 ms to prevent carryover of the beat from previous sequences (e.g., the next trial started between 325 and 375 ms after the last beat in the silence, which is not on the beat, nor at a subdivision of the beat). On trials that contained a probe tone, we chose to only use the last three probe positions, to 1) limit the time for the EEG experiment to prevent fatigue, 2) still provide participants with the incentive to form expectations well into the silence period, as a probe tone could appear as late as 1485 ms into the trial, and 3) obtain some measure of whether participants formed expectations, by including positions that were expected based on the beat and the pattern (1200 ms), based on the pattern only (990 ms) or neither (1485 ms).

### Procedure

Participants were tested individually in a dedicated lab at the University of Amsterdam. Upon arrival, participants were provided with information about the experiment, provided written informed consent, and were allowed to practice the task. On probe tone trials (all trials in Experiment 1, 25% of the trials in Experiment 2), participants were asked to judge on a four-point scale (“very poorly”, “poorly”, “well”, “very well”) how well the probe tone fitted the preceding sequence, similar to previous studies investigating the perception of musical meter (Manning et al., 2017; Manning & Schutz, 2013; Palmer & Krumhansl, 1990). Participants could respond with four buttons on the armrest of their chair, two on each side. The order of the answer options on the screen in front of the participants was randomized on each trial, to avoid any artefacts of motor preparation in participants that anticipated which answer they would provide. There was no time limit for responses and the next trial started after a response was made.

In Experiment 1, each participant was presented with 18 blocks of 30 trials, amounting to 540 trials in total (30 trials per probe position for each condition). In Experiment 2, participants were presented with 18 blocks of 36 trials, for a total of 648 trials (216 per condition). For each condition, 162 trials were silence trials, and did not contain a probe tone, and 54 trials contained a probe tone (18 per position). The number of probe trials per block was varied between 3 and 11. In both experiments, in each block, only one type of sequence (beat-based, pattern-based, or random) could appear to optimize for the formation of expectations. Blocks were semi-randomized, with each type appearing once in a set of three blocks.

Sounds were presented through a single speaker positioned in front of the participant using Presentation® software (version 14.9, www.neurobs.com). After the experiment, participants filled out the musical training subscale from the Goldsmith Musical Sophistication Index (GMSI) questionnaire (Müllensiefen et al., 2014). In total, a behavioral session lasted two hours, and the EEG session lasted between 3.5 and 4 hours.

### Behavioral analysis

A total of 17280 responses was included in the analysis of Experiment 1 (32 participants, 3 conditions, 6 probe positions, 30 responses each), and 4374 in the analysis of Experiment 2 (27 participants, 3 conditions, 3 probe positions, 18 responses each). To account for the ordinal nature of the Likert-scale responses (Bouwer et al., 2018; Carifio & Perla, 2008; Jamieson, 2004), we used a mixed ordinal regression model. With this model, the ordinal responses are normalized, to correct for potential unequal distances between rating points. The results can subsequently be interpreted similar to the results from a normal mixed model regression. Two independent variables were included in the model as fixed factors: Condition (beat-based, pattern-based, or random), and Probe Position (600, 780, 885, 990, 1200, or 1485 ms; only the latter three in Experiment 2). Additionally, GMSI score was included as a continuous variable (Musical Training), as well as interactions between all factors, and a random intercept for each subject to account for between-subject variation.

The initial model showed a significant effect of Probe Position in the random condition, most likely due to recency effects. To assess the effect of Probe Position in the beat-based and pattern-based conditions while accounting for this, for each participant we subtracted the mean response in the random condition at each position from the responses in the beat-based and pattern-based condition. Subsequently, we used these baseline-corrected ratings in a second ordinal regression model, with only two levels for the factor Condition (beat-based and pattern-based) and without the random intercept for each participant (as the baseline correction already corrected for between-subject variability). For both the original model, and the baseline corrected model, significant interactions were followed up by tests of simple effects, corrected for multiple comparisons using a Bonferroni correction.

The statistical analysis was conducted in R (R Development Core Team, 2008). The ordinal mixed model was implemented using the clmm() function from the ordinal package (Christensen, 2019). Subsequently, we used the Anova() function from the car package (Fox & Weisberg, 2019) to look at omnibus effects for the main factors of interest, and the emmeans package (Lenth, 2019) to assess simple effects and compare slopes between conditions.

### EEG recording

EEG was recorded at 1024 Hz using a 64-channel Biosemi Active-Two acquisition system (Biosemi, Amsterdam, The Netherlands), with a standard 10/20 configuration and additional electrodes for EOG channels (located under and on the left and right sides of the eye), on the nose, on both mastoids, and on both earlobes.

### EEG analysis

Preprocessing was performed in MATLAB, version 2015a (Mathworks) and EEGLAB, version 14.1.1 (Delorme & Makeig, 2004). Data were offline down-sampled to 256 Hz, re-referenced to averaged mastoids, bad channels were manually removed, and eye-blinks were removed using independent component analysis. Subsequently, bad channels were replaced by values interpolated from the surrounding channels.

#### ERPs

For the ERP analysis, the continuous data were filtered using 0.1 Hz high-pass and 40 Hz low-pass finite impulse response filters (as implemented in the standard EEGLAB filter function pop_eegfiltnew). Epochs were extracted from the data from -1800 till 2100 ms relative to the onset of the last sound. Epochs with a voltage change of more than 150 microvolts in a 200 ms sliding window were rejected from further analysis. For each participant and condition, epochs were averaged to obtain the ERPs, and ERPs were averaged over participants to obtain grand average waveforms for plotting. All waveforms were initially baseline corrected using the average voltage of a 50 ms window preceding the onset of the last sound of the sequence. This baseline can be regarded as preceding the “cue” (the last event before the onset of the expectation). Such a baseline is customary in CNV analyses. However, visual inspection suggested that this baseline was biased, as baseline correction resulted in an overall shift of the waveform amplitude relative to each other, as also reflected in a significant cluster when comparing beat-based and pattern-based conditions that spanned the entire analysis epoch (see Appendix, Figure A1). This was likely caused by the rapid succession of sounds preceding the onset of the silence, which made it impossible to find a clean, unbiased baseline. Therefore, we repeated the analysis without baseline correction, to confirm that the results were not caused by a noisy baseline.

As a control analysis, we repeated the above analysis for the longest intervals during the presentation of the rhythmic streams, to assess whether in those longer intervals, we could observe similar deflections in the evoked potentials as observed in the silence. For this analysis, epochs were extracted from -200 till 880 ms around the onset of the sound preceding the long interval (600 ms in the beat-based condition, and 780 ms in the other conditions, see Figure 1). All preprocessing steps were identical to the analysis of the ERPs in the silence. The control analysis is reported in the Appendix (Figure A1).

Three cluster-based permutation tests (Oostenveld et al., 2011) were used to compare all three conditions against each other (i.e., beat-based vs. random; pattern-based vs. random; beat-based vs. pattern-based), comparing all timepoints from 300 till 1200 ms after the onset of the final sound for the silence (see Figure 1), and all timepoints from 300 till 600 ms after the onset of the preceding sound for the long intervals during the sequences (as the next sound came in at 600 ms for the beat-based condition, we could not compare the conditions beyond this timepoint). This window excluded a large portion of the ERP response to the previous sound, and included both the first expected moments for beat-based (600 ms) and pattern-based (780 ms) expectations in the silence window, and additional time to allow for an evaluation of possible return to baseline of the CNV (Breska & Deouell, 2017). For the ERP analysis, clusters were formed based on adjacent time-electrode samples.

For all EEG analyses, cluster-based tests were evaluated statistically by forming clusters of samples based on dependent samples t-tests and a threshold of *p* < 0.05, and using permutation tests with 2000 permutations of the data. We report corrected *p-*values to account for two-sided testing (multiplied by a factor of two).

#### Frequency-domain analysis

To obtain the spectrum of the EEG signal in the silence, we used the raw, unfiltered data. Epochs were extracted from the continuous data both from - 1800 till 0 ms relative to the onset of the last sound (control window, see Figure 1), and from 300 till 2100 ms relative to the onset of the last sound (silence window, see Figure 1), the latter starting at 300 ms to avoid contamination from the final ERPs. Both windows thus had equal length, both spanning three full cycles of the beat. Epochs with an amplitude change of 250 microvolts or more in a sliding 200 ms window were rejected from further analysis. The more lenient rejection criterium compared to the ERP analysis was used to account for the fact that these data were unfiltered, and to avoid rejection of too many trials that showed some slow drift. All epochs were baseline corrected using the mean of the entire epoch. Subsequently, epochs were averaged for each condition separately to obtain the evoked signal, phase locked to the onset of the final sound, and similar to previous studies using frequency tagging to look at beat-based perception (Lenc et al., 2021; Nozaradan et al., 2011, 2012).

For each participant and condition separately, the average waveforms were transformed into the frequency domain using an FFT, with the data zero-padded to 4608 samples (NFFT) to obtain a better frequency resolution (0.056 Hz), and importantly, be able to extract data at exactly the frequencies of interest. Note that the zero-padding can only improve the frequency resolution, but not the frequency precision, which by definition with the 1800 ms epochs is limited to 0.56 Hz. While the design of the experiment simply does not allow for a better resolution, the 0.56 Hz does allow us to differentiate between the frequencies of interest, which are 0.56 Hz or more apart. The obtained power values at each frequency were normalized to account for the 1/f distribution of noise (Nozaradan et al., 2011, 2012), by subtracting the average of neighboring bins four to six on either side for all frequencies (e.g., 1.33 – 1.44 Hz and 1.89 – 2.00 Hz for the beat frequency, 1.89 – 2.00 Hz and 2.44 – 2.56 Hz for the pattern-based frequency, and 3.00 – 3.11 Hz and 3.56 – 3.67 Hz for the beat subdivisions). To account for bleeding into neighboring frequency bins (Nozaradan et al., 2011, 2012), for each frequency, we averaged over 5 bins centered on that frequency (e.g., for the frequencies of interest, this was 1.56 – 1.78 Hz for the beat frequency, 2.11 – 2.33 Hz for the pattern-based frequency, and 3.22 – 3.44 Hz for the beat subdivisions).

To statistically test differences between conditions in the evoked power at the frequencies of interest, we used cluster-based permutation tests. First, this avoided bias by selecting only a subset of electrodes, as we used all scalp electrodes, as was done in previous research (Lenc et al., 2018; Nozaradan et al., 2011; Tal et al., 2017). Second, the permutation tests accounted for the non-normal distribution of the data. Like for the ERPs, we ran t-tests comparing the normalized data for all conditions and for each frequency of interest (e.g., those most prominent in the sound signal). We included the frequencies that showed the highest peaks in the spectral analysis of the sound (i.e. 1.67 Hz, 2.22 Hz, and 3.33 Hz), except for 3.89 Hz, since a peak at this frequency was absent on visual inspection in the spectral decomposition of the EEG data. For the frequency-domain analysis, clusters were formed based on adjacent electrodes.

The cluster-based tests yielded null results for the pattern-based condition in the silence (e.g., there was no larger power at 2.22 Hz in the pattern-based than in the random condition, see Results). To quantify the possible absence of persistent entrainment for the pattern-based condition, we performed a Bayesian t test using JASP (JASP, 2019; Wagenmakers et al., 2018). We compared the power in the pattern-based and random conditions at 2.22 Hz in the silence, averaged over electrodes contributing to the significant cluster in the silence for the beat-based condition at 1.67 Hz, to optimize for finding entrainment effects. We estimated Bayes factors using a Cauchy prior distribution (r =.71) and performed a robustness check to assess the effect of a different prior (r = 1; see (Jeffreys, 1961; Wagenmakers et al., 2018)).

#### Multiscale entropy (MSE)

MSE is a measure of signal irregularity. To compute MSE, the EEG signal is divided into patterns of a certain length, and throughout the signal, the number of repeating patterns is counted. More repetitions indicate a more regular signal, and yield a lower entropy value. By calculating entropy for patterns of different lengths (“multiscale”), the contributions of slower and faster timescales in the signal can be assessed. However, the mapping between entropy timescales and spectral frequencies is not absolute, especially since entropy is not per se related to a signal being oscillatory in nature (Kloosterman et al., 2020; Kosciessa et al., 2020). The advantage of using MSE is that it does not require filtering of the data, and it does not assume stationarity (e.g., it can pick up on regularities that are asymmetrical, or that do not have a fixed amplitude or period). Here, we computed MSE on the control and silence epochs separately.

We computed MSE on high-pass filtered data (0.5 Hz). Epochs were extracted identical to the epochs for the frequency domain analysis. To compute MSE, we used the mMSE toolbox, a plugin to the Fieldtrip toolbox (Kloosterman et al., 2020), with *m* = 2 and *r* = 0.5, as was done previously for EEG data (Kloosterman et al., 2020). For details on how MSE is computed, see the Appendix. As for the frequency-domain analysis, we used cluster-based permutation tests to assess statistical significance. For each comparison between conditions, we used paired t-tests comparing each electrode-timescale combination (note that the above computation of entropy yields one value per condition for each electrode and timescale) to form clusters.

#### Multivariate decoding

Our exploratory decoding approach is based on the assumption that temporal expectations are always coupled with feature or spatial expectations (e.g., we cannot predict “when” without also predicting “what”), as suggested by studies showing that we only use temporal expectations to improve perception if we can also predict the content (Morillon et al., 2016; Wollman & Morillon, 2018) or location (O’Reilly et al., 2008) of an upcoming event. Thus, we expected to be able to decode the representation of the expected sound at expected moments. As the expected moments are different for each condition, this then allows us to decode in the silence window whether participants were previously listening to a beat-based, pattern-based, or random sequence.

The decoding was conducted on data that was preprocessed in a similar way as for the MSE analysis, but with epochs extending from -1800 to 2100 ms relative to the onset of the last sound. Since the decoding is done in a time resolved way (e.g., sample by sample), there is no need to leave out the response to the ERPs in the analysis. Additionally, the data were resampled to 32 Hz to increase signal to noise, using shape-preserving piecewise cubic interpolation, as implemented in the Fieldtrip toolbox (Oostenveld et al., 2011). Using the ADAM toolbox (Fahrenfort et al., 2018), we applied a classification algorithm to the preprocessed data for each participant. Using all electrodes, each dataset was split into 10 equally sized subsets, for 10-fold cross-validation of the decoding. For each subset, a linear discriminant classifier trained on the remaining 9 subsets was tested. Each condition was decoded against both other conditions (e.g., beat-based vs. pattern-based, beat-based vs. random, and pattern-based vs. random), creating a temporal generalization matrix of classification accuracy at each possible combination of training and testing time points (King & Dehaene, 2014). Subsequently, we examined whether we could observe a pattern of recurrent activity (King & Dehaene, 2014). Classification accuracies averaged over the 10 folds for each comparison of two conditions for the silence window (300-2100 ms after the onset of the last sound) were submitted to cluster-based permutation tests to assess whether they exceeded the chance level of 0.5. Clusters were based on T-tests with a threshold of 0.05 for each training-testing time point combination, comparing the accuracy to 0.5.

The initial decoding analysis yielded large effects that may be task-related (see Results for an explanation of these results). Therefore, we ran an additional decoding analysis in which we decoded expected and unexpected positions against each other within each condition. Details on this additional analysis can be found in the Appendix.

## Results

### Behavioral effects of beat-based expectations last multiple beat cycles, while those of pattern-based expectations reflect one interval

Figure 3A shows the average ratings for each condition and probe position from Experiment 1. Visual inspection of this figure suggests that beat-based expectations were associated with higher fitness ratings for sounds at expected times than unexpected times for two beat cycles in the silence window (at 600 and 1200 ms), while the effects of pattern-based expectations appeared to reflect mainly the first expected time point in the silence window (780 ms). This was confirmed by our statistical analyses. The ordinal regression showed main effects of Condition (*χ*^2^(2) = 300.72, *p* < 0.001), Position (*χ*^2^(5) = 1067.05, *p* < 0.001), as well as Musical Training (*χ*^2^(1) = 5.16, *p* = 0.023). However, crucially, these main effects were accompanied by a very large two-way interaction between Condition and Position (*χ*^2^(10) = 2478.98, *p* < 0.001), showing that the effects of beat-based and pattern-based expectations on fitness ratings differed, depending on the position of the probe. We found additional smaller interactions between Position and Musical Training (*χ*^2^(5) = 67.26, *p* < 0.001), Condition and Musical Training (*χ*^2^(2) = 6.01, *p* = 0.05), and, interestingly, Condition, Position, and Musical Training (*χ*^2^(10) = 204.88, *p* < 0.001). Following the interactions, tests of simple main effects showed that the effect of Position was significant in all conditions (all *p*s < 0.001), and the effect of Condition was significant for all probe positions (all *p*s < 0.001). The main effect of Position in the random condition showed that even after sequences in which no specific temporal structure was present, ratings depended on the position of the probe. This likely was due to recency effects. To account for these effects when comparing the ratings at different positions in the beat-based and pattern-based conditions, we subtracted the ratings in the random condition at each position from the ratings in the other conditions. The baseline corrected model (Figure 3B) showed similar interactions between Position and Condition (*χ*^2^(5) = 2147.55, *p* < 0.001), and between Position, Condition, and Musical Training (*χ*^2^(5) = 154.46, *p* < 0.001). The latter indicated that the correlation between Musical Training and the rating score depended on both the Position and the Condition of the probe tone.

**Figure 3.**
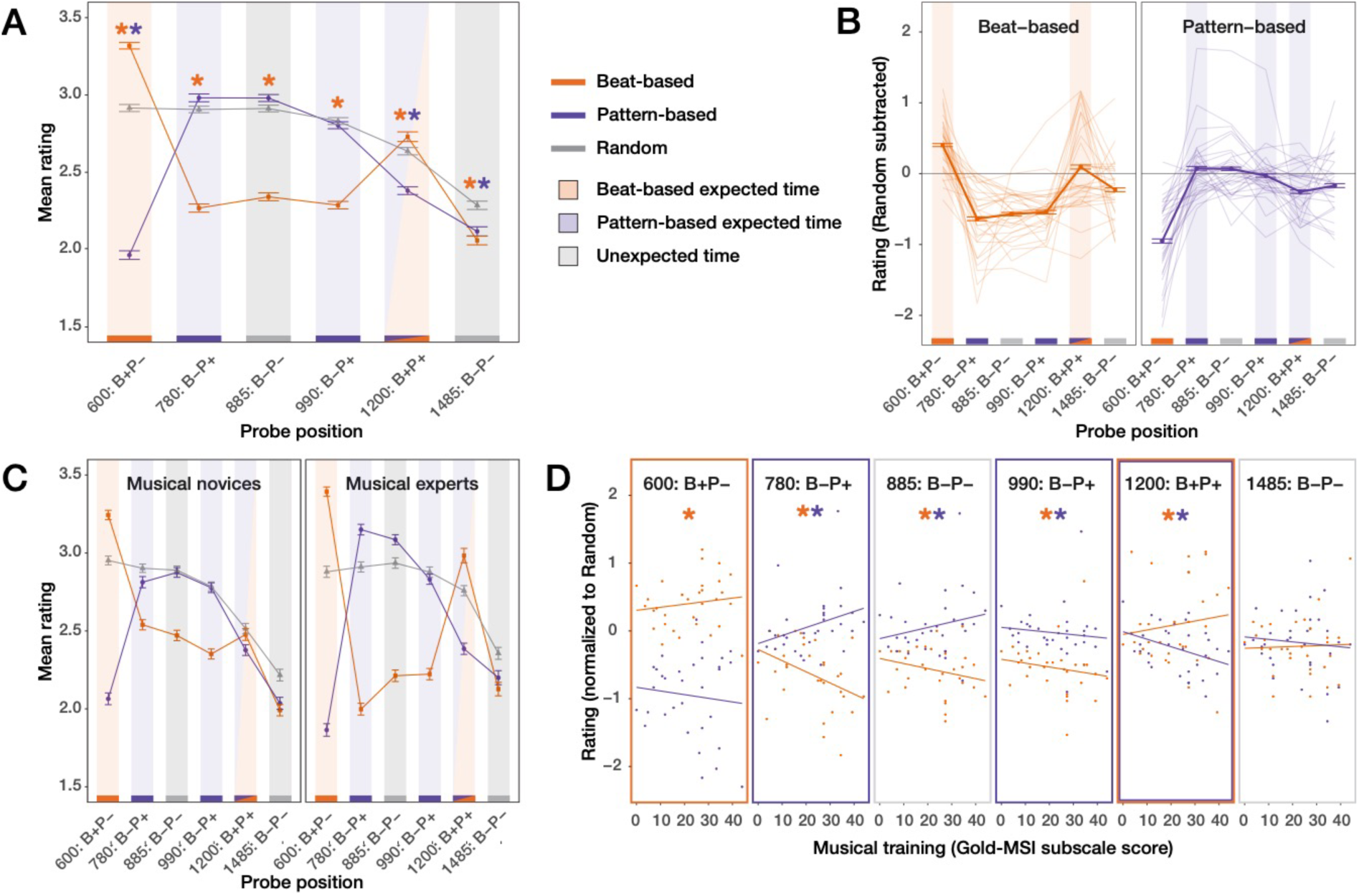
The effects of beat-based expectations on fitness ratings can be differentiated from those of pattern-based expectations, and are associated with musical training. A) Mean ratings for all conditions and positions. Colored asterixis indicate positions where ratings in the beat-based (orange) and pattern-based (purple) conditions differed from the random condition at *p* < 0.05 (Bonferroni corrected). B) Single participant data, with the random condition subtracted to account for serial position effects. The expectedness pattern is indicated by colored lines on the bottom of the plots (orange: expected based on the beat; purple: expected based on pattern; grey: neither). For the beat-based condition, ratings followed this pattern for two beat cycles. For the pattern-based condition, ratings followed the pattern for one interval. C) Data median split based on scores on the musical training questionnaire. The pattern of results, while present for both groups of participants, is enhanced for the group of participants with most musical training (“experts”). Note: the median split is for visualization purposes only, the models were run with musical training as a covariate. Error bars in panels A-C are 2 standard errors (note: these are computed on the complete dataset, not the participant averages, as the ordinal model is run on trial-level data). D) Association between musical training and rating for each condition and position. Colored asterixis show positions in which the association between musical training and the ratings was significantly correlated (*p* < 0.05, Bonferroni corrected). A positive association was observed for the beat-based condition at 600 and 1200 ms (expected positions) and for the pattern-based condition at 780 (expected) and 885 ms (unexpected). Negative associations were observed in the beat-based condition at 780, 885, and 990 ms (all unexpected), and in the pattern-based condition at 990 (unexpected) and 1200 ms (expected).

#### Beat-based expectations

At both 600 ms and 1200 ms (expected in terms of a beat), probes in the beat-based condition were rated as better fitting than probes in the random condition (*p* < 0.001 and *p* = 0.004) as evident from the full model; all simple effects from the full model can be found in the Appendix, Table A1). At 780, 885, 990, and 1485 ms (unexpected in terms of a beat), probes in the beat-condition were rated as worse fitting than probes in the random condition (all *p*s < 0.001). Moreover, within the beat-based condition, at 600 ms, baseline corrected ratings were higher than at any other probe position (all *p*s < 0.001), and at 1200 ms baseline corrected ratings were higher than at 780, 885, 990, or 1485 ms (all *p*s < 0.001). Baseline corrected ratings for probes at 780, 885, and 990 ms (all unexpected in terms of the beat) did not differ from each (all *p*s > 0.93). Probes at 1485 ms (unexpected in terms of the beat) were rated as better fitting than probes at 780, 885, and 990 ms (all *p*s < 0.001). All simple effects from the corrected model can be found in the Appendix, Table A2.

As can be seen in Figure 3C and 3D, higher scores on the Musical Training questionnaire were associated with higher fitness ratings in the beat-based condition at 600 and 1200 ms (expected in terms of the beat), but lower fitness ratings at 780, 885, 990, and 1485 ms (unexpected in terms of the beat). In other words, musically trained participants were better able to differentiate between probes that were in expected and unexpected positions (Figure 3C). Slopes reach significance at all positions except 1485 ms (all *p*s < 0.022). Also, the association between Musical Training and ratings differed between beat-based and pattern-based conditions, at 600, 780, 885, and 1200 ms (all *p*s < 0.004).

To sum up, for the beat-based sequences, we could observe a clear pattern in the results indicating that beat-based expectations were used to rate the probes well into the silence window, affecting ratings up to 1200 ms after the onset of the last sound. Beat-based expectations lead to higher ratings for expected probes (600 and 1200 ms), and lower ratings for unexpected probes (780, 885, 990, and 1485 ms), both when comparing ratings for each position to the random condition, and when comparing ratings for each position within the beat-based condition. At 1200 ms, these effects resulted in a classic inverted U-curve, as previously associated with beat-based processing (Bauer et al., 2015; Jones et al., 2002), with optimal performance on the beat, and diminished performance on either side (e.g., both earlier and later). The effects of beat-based expectations did diminish over time, as is apparent from differences between ratings at 600 and 1200 ms, and at 1485 ms and other unexpected time points. Both the enhancing and attenuating effects of beat-based expectations were correlated with musical training. It is worth noting that the longer lasting effects of beat-based expectations (at 1200 ms) were very heterogenous in our participant pool. Out of 32 participants in Experiment 1, only 18 showed the inverted U, with higher ratings at 1200 than at 990 and 1485 ms.

#### Pattern-based expectations

For pattern-based sequences, ratings at 600 ms (unexpected based on the pattern) were lower than for the random sequences (*p* < 0.001) and lower than at any other position (all *p*s < 0.001, baseline corrected model), showing that participants also formed predictions based on the sequences. In line with this, at 780 ms (expected in terms of the pattern), ratings were numerically higher in the pattern-based condition than in the random condition, though this difference did not survive the Bonferroni correction (*p* = 0.058). After this point, ratings did not differ between pattern-based and random conditions for probes at 885 (unexpected) and 990 (expected) ms, suggesting that the responses followed the rhythmic pattern mainly in the beginning of the silent period. In line with this, in the remainder of the silence window, the ratings continued to deviate from what would be predicted based on the pattern, with lower ratings for the pattern-based than random condition at 1200 ms (expected; *p* < 0.001). Ratings were also lower for the pattern-based than random condition at 1485 ms (unexpected, *p* < 0.001), and ratings at 780 ms (expected) did not differ from ratings at 885 ms (unexpected), while being marginally higher than at 990 (expected, *p* = 0.080), and higher than at 1200 (expected), and 1485 (unexpected) ms (both *p*s < 0.032). In addition, ratings at 885 and 990 ms were higher than at 1200 and 1485 ms (all *p*s < 0.002). See the Appendix, Tables A1 and A2, for all simple effects.

As for beat-based expectations, for pattern-based expectations, there was a positive association between ratings and musical training at an expected time point (780 ms; *p* < 0.001), and a negative, albeit nonsignificant, association at an unexpected time point (600 ms). Thus, like for beat-based expectations, at these early time points, musicians were better able than non-musicians at differentiating between expected and unexpected moments in time. However, at 885 ms (unexpected in terms of the pattern), the results behaved like at 780 ms, with higher ratings associated with more Musical Training (*p* = 0.002). At 990 and 1200 ms, Musical Training was associated with lower ratings (both *p*s < 0.05), but these results are counterintuitive, as these are expected positions based on the pattern.

The results for pattern-based expectations suggest that just like for beat-based expectations, participants were able to predict the timing of probes based on the preceding sequence. However, the results show that while this was still the case at 780 ms after the onset of the last tone, at later probe positions, the effects of pattern-based expectations did not reflect the preceding sequence. At 885 ms, the results, both in terms of the ratings and how they were associated with musical training, behaved similar to at 780 ms. After this point, the results suggest that participants did not use the preceding sequence to guide their responses, but instead, used a different heuristic.

Figure 4 shows the behavioral results obtained from the EEG experiment. Replicating Experiment 1, we found main effects of Condition (*χ*^2^(2) = 92.30, *p* < 0.001) and Position (*χ*^2^(2) = 49.36, *p* < 0.001), accompanied by interactions between Condition and Position (*χ*^2^(4) = 96.62, *p* < 0.001), and Condition, Position, and Musical Training (*χ*^2^(4) = 11.04, *p* = 0.03). Following the analysis strategy from Experiment 1, we subtracted the ratings from the Random condition from the ratings for the other two conditions, yielding a baseline corrected model with similar interactions (Condition and Position: *χ*^2^(2) = 31.46, *p* < 0.001; Condition, Position, and Musical Training: *χ*^2^(2) = 8.29, *p* = 0.02).

**Figure 4.**
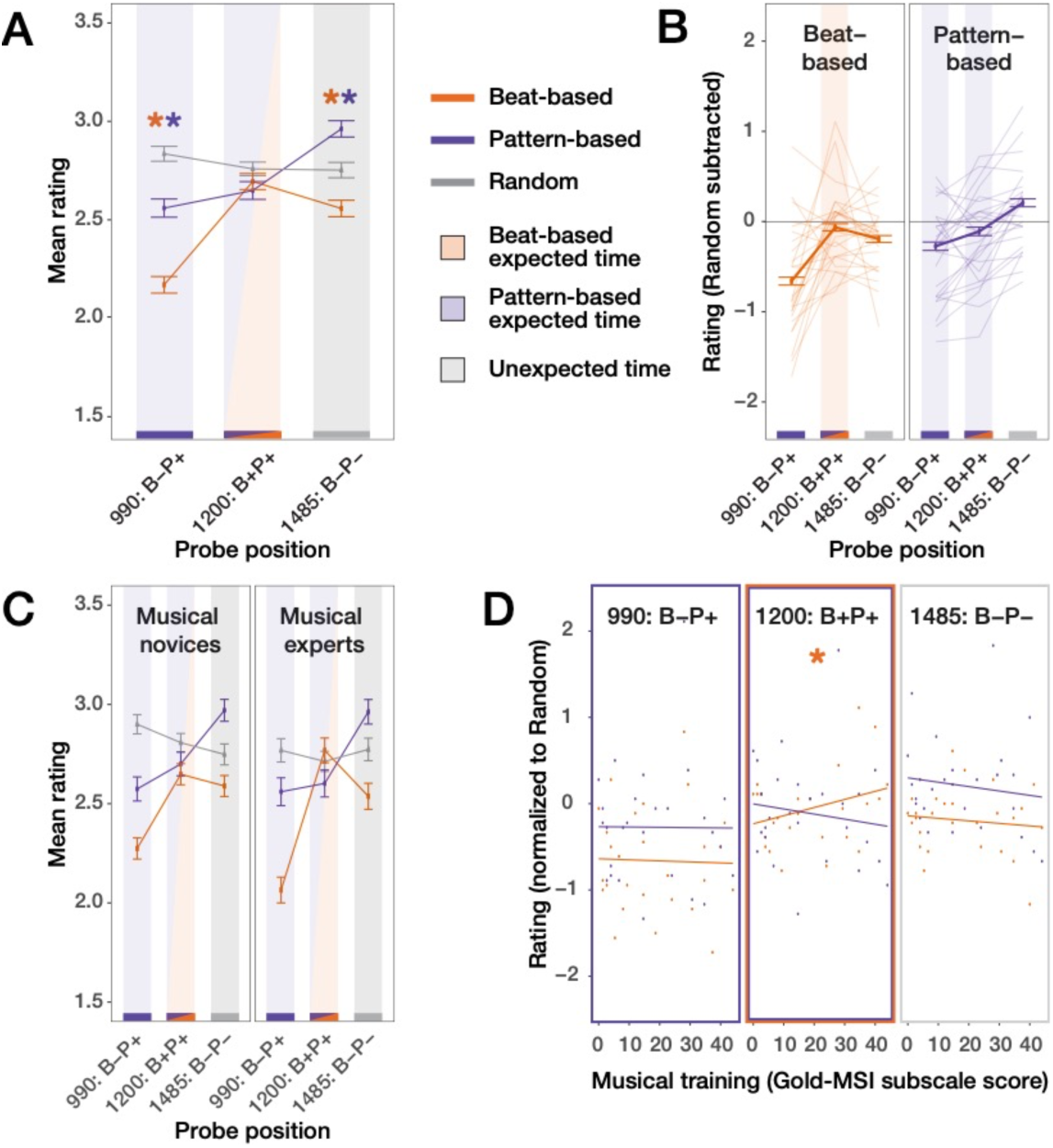
Behavioral results during the EEG experiment replicate findings from Experiment 1. Even with only 18 trials per participant per condition and position, we could replicate the inverted U-curve for beat-based sequences at the end of the silence epoch. Like in Experiment 1, the results for the pattern-based condition do not follow the pattern, but instead, are consistent with building expectations for the next trial. A) Mean ratings for all conditions and positions. Colored asterixis indicate positions where ratings in the beat-based (orange) and pattern-based (purple) conditions differed from the random condition (with *p* < 0.05, Bonferroni corrected). B) Single participant data, with the random condition subtracted to account for serial position effects. C) Data median split based on scores on the musical training questionnaire. Error bars in panels A-C are 2 standard errors. D) Association between musical training and rating for each condition and position. See Figure 3 for more details.

In line with the preceding beat-based sequences, probes at 990 and 1485 ms (both unexpected times based on the beat) were rated lower in the beat-based condition than in the random and pattern-based conditions (all *p*s < 0.012), and within the beat-based condition, probes at 1200 ms were rated as better fitting than at 990 (*p* < 0.001) and 1485 ms, though the latter difference did not reach significance. Thus, like in Experiment 1, we found an inverted U-curve at 1200 ms after the final tone, suggestive of beat-based expectations lasting at least two beat cycles. As in Experiment 1, at 1200 ms (expected based on the beat), higher ratings in the beat-based condition were associated with more musical training (*p* = 0.047), suggesting that the effects of beat-based expectations correlate with musical expertise. Additionally, probes at 990 ms were rated lower than at 1485 ms (*p* < 0.001), possibly because the effects of beat-based expectations diminished over the course of the silence.

In the pattern-based condition, ratings did not follow the pattern of the preceding sequences. At 990 ms (expected based on the pattern), probes were rated as worse fitting than in the random condition (*p* < 0.001), and as worse fitting than at 1200 (expected) and 1485 (unexpected) ms (both *p*s < 0.023), while at 1485 ms (unexpected), probes were rated as better fitting than in the random condition, and as better fitting than at 1200 (expected) ms (both *p*s < 0.001). Also, at 1200 ms (expected), higher ratings were associated with less musical training for the pattern-based condition (though after the Bonferroni correction, only marginally so: *p* = 0.06), contrary to what would be expected if the effects of expectations are enlarged in musical experts. All simple effects can be found in the Appendix, Tables A1 and A2.

Thus, the behavioral results from Experiment 2, though based on less trials than Experiment 1, suggest a similar pattern as found in Experiment 1: while beat-based expectations exert their effect well into the silent period, with participant faithfully following the beat in their goodness-of-fit ratings, pattern-based expectations do not affect ratings in the second half of the silent period in a manner consistent with the learned pattern. Albeit speculatively, the results for the pattern-based expectations may be more in line with expectations for the start of the next trial leading to higher ratings for probe positions closer to the end of the silent period, as in Experiment 2, after non-probe trials, the next trial followed each silent period at a somewhat predictable time.

### Differences in the evoked potential elicited by beat-based and pattern-based sequences

Figure 5 shows the average ERPs for each condition, without baseline correction (see Materials and Methods), and scalp topographies for windows in which we found significant clusters. In line with previous research, we may expect climbing neuronal activity, or a CNV, flexibly adapting its slope to peak at the moment that participants expect the next event (Breska & Deouell, 2017; Breska & Ivry, 2020; Damsma et al., 2021; Mento, 2013). We did observe differences in the ERPs between conditions, but the peak of the differences was not at the expected time, but rather, fell earlier (see the Appendix, Figure A1, for the difference waveforms between conditions). Without baseline correction, the random condition elicited a significantly more negative ERP (*p* = 0.011) than the beat-based condition in a frontocentral cluster between 300 and 614 ms after the onset of the last sound (though note that we did not include timepoints preceding 300 ms in the cluster-based tests). Likewise, in the same latency range (300 – 473 ms), there was a trend for the pattern-based condition to elicit a more negative ERP than the beat-based condition (*p* = 0.077). Thus, in a window between approximately 300 and 450 ms, we found tentative evidence for more negative-going waveforms in both the random and pattern-based condition compared to the beat-based condition, with a central scalp topography (Figure 5B). In a later window, the pattern-based condition elicited a second negative deflection, which showed a trend to be larger than in the beat-based condition (*p* = 0.077, 864 – 1005 ms). While these results were somewhat different depending on the choice of baseline (see Figure A1 in the Appendix for the results with a traditional pre-cue baseline), the overall picture is the same, with significant clusters in an early and later window indicating differences between conditions in the ERPs.

**Figure 5.**
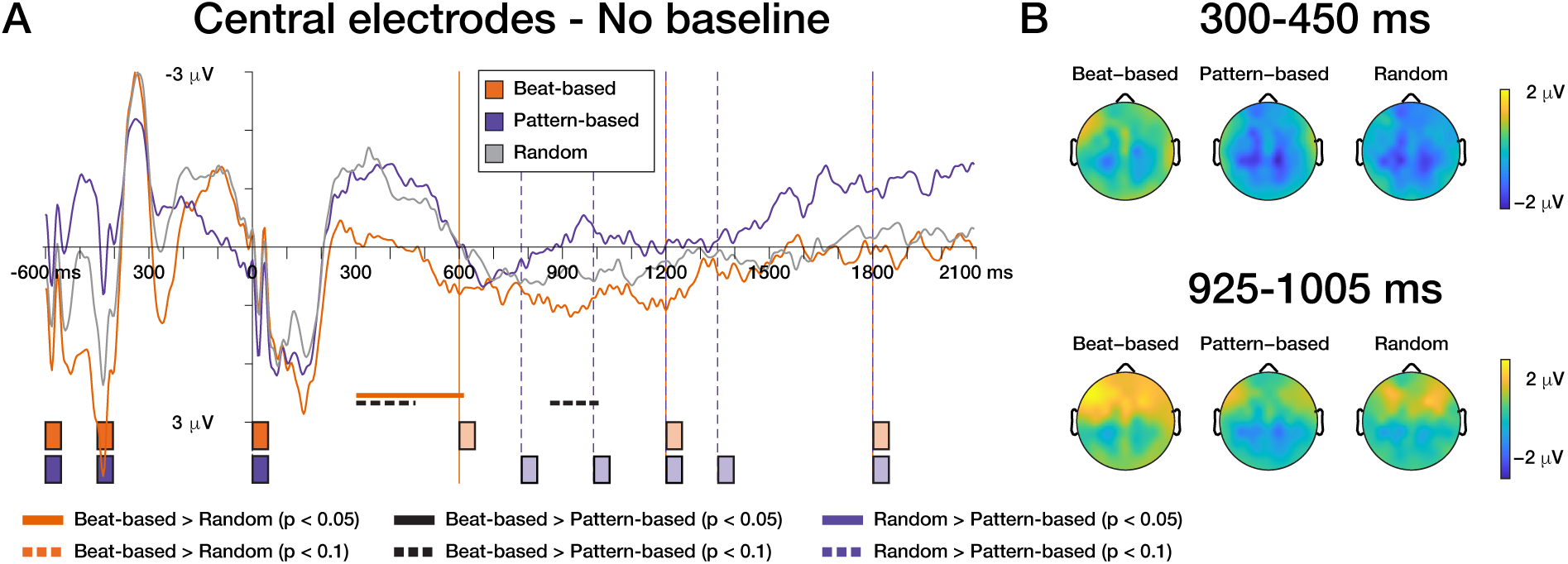
Beat-based and pattern-based can be differentiated based on ERPs. A) Left panel show the grand average waveforms for the silence window for a central electrode cluster (FC1, FCz, FC2, C1, Cz, C2, CP1, CPz, CP2). Time 0 is the onset of the last tone of the sequence. Colored bars on the bottom of the plots, and vertical orange and purple lines, indicate at which times a tone would be expected based on the beat (light orange) and the pattern (light purple). Note that these are expected times, but no sounds were played during the window shown after time 0. Sounds during the sequence are depicted in dark orange and purple. B) The scalp distributions for windows in which a significant cluster was observed.

The time course of the effect deviated from what we expected: the negative deflection did not peak at the next expected moment in time, but rather, peaked much earlier. Also, contrary to previous research (Breska & Deouell, 2017), the beat-based condition elicited the most positive-going ERP, instead of a typical CNV. To further explore and confirm these results, as a control analysis, we performed the same ERP analysis on the longest time intervals during the sound presentation. The waveforms showed a negative deflection very similar in morphology and scalp distribution to the one we found in the silence window (see Figure A1 in the Appendix). During the sound presentation, like in the silence, this negative deflection was largest for the pattern-based condition, though the difference was only significant when comparing the pattern-based condition to the random (*p* = 0.039) but not the beat-based condition (*p* = 0.1). The latter non-significant result may be due to a noisy baseline, as during sound presentation, the succession of intervals in the beat-based and random sequences was (semi-)randomly chosen, and therefore, the baseline was not consistent over conditions. Overall, however, this control analysis yielded very similar results to the analysis of ERPs in the silence window.

### Frequency-domain analysis shows persistent power at the beat frequency following beat-based sequences

A strong prediction of entrainment theories is that the entrainment outlasts stimulation with the entraining stimulus. Therefore, we next looked at the frequency content of the EEG signal in the silent period. Specifically, we predicted that if entrainment occurs, we would find enhanced power at the frequencies associated with the rhythmic sounds to be found in the silence. In Figure 6A, the average power for all electrodes, separated for each condition in the control (i.e. during the auditory sequence) and silence windows is depicted as a function of frequency. In the control window (Figure 6, left, and Figure 7, top), the frequency response followed the sound input. That is, at the beat frequency (1.67 Hz), significant clusters indicated higher power in the control window for the beat-based sequences than the pattern-based (*p* < 0.001) and random (*p* = 0.006) sequences. At 2.22 Hz, prominent in the pattern-based and random sound sequences, higher power was observed in the EEG signal in the pattern-based than beat-based (*p* = 0.02) and random (*p* = 0.023) sequences, and higher power was observed in the random than beat-based sequences (*p* = 0.035). Finally, at 3.33 Hz (subdivisions of the beat), power in the control window was larger for the beat-based than pattern-based (*p* = 0.013) and random (*p* = 0.005) conditions. Thus, in the control window, the EEG signal reflected the spectral properties of the sound signal, which is likely caused by each sound eliciting an ERP, which are represented in steady-state potentials, and thus picked up by the frequency analysis (Keitel et al., 2021), especially since the ERPs elicited by our stimuli extended for hundreds of milliseconds, and our analysis focused on frequencies >1 Hz (Zhou et al., 2016).

**Figure 6.**
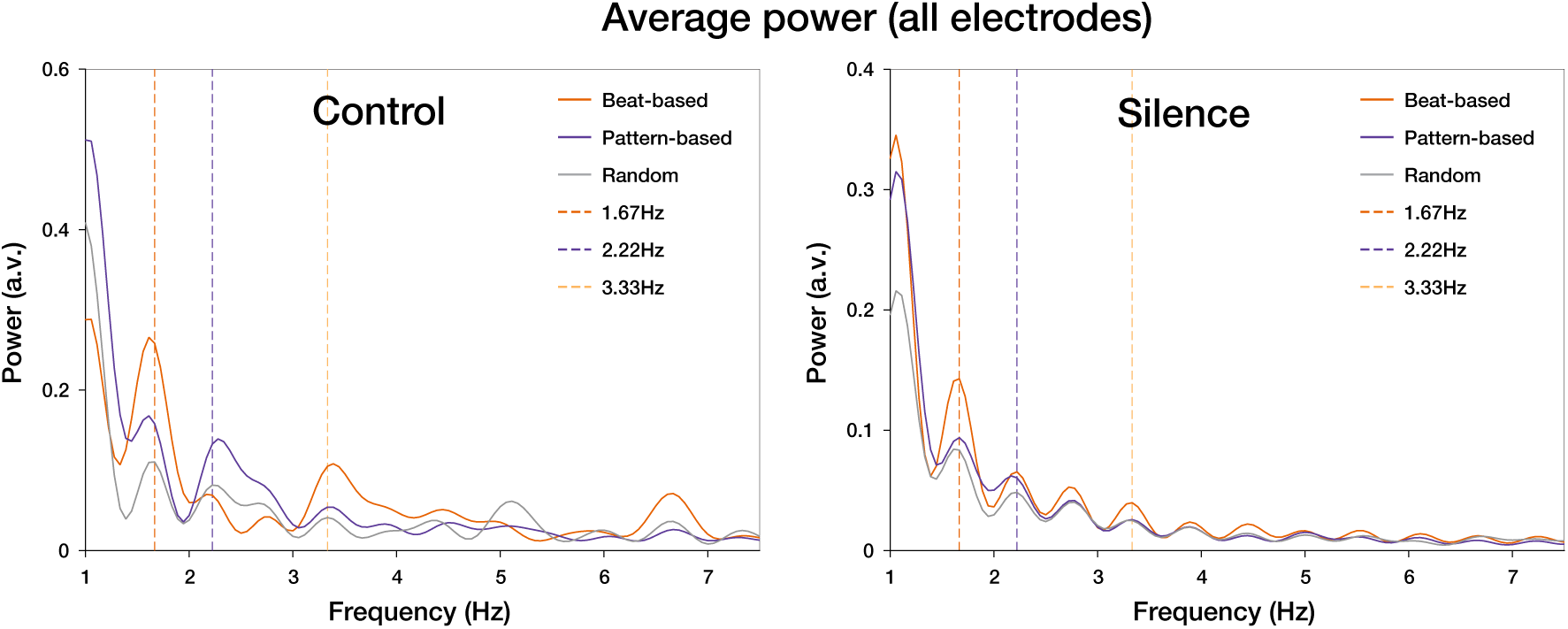
Power at the beat frequency persists during the silence window. In the control window, during auditory stimulation, peaks in power can be observed at all frequencies of interest, and for the relevant conditions (1.67 and 3.33 Hz in the beat-based condition, and 2.22 Hz in the pattern-based condition). In the silence window, the peaks in power at 1.67 and 3.33 Hz were larger in the beat-based condition than in the pattern-based and random conditions, while peaks at 2.22 Hz did not differ between conditions. Note: the raw data is depicted here, before the normalization procedure. Data shown is averaged over all electrodes.

**Figure 7.**
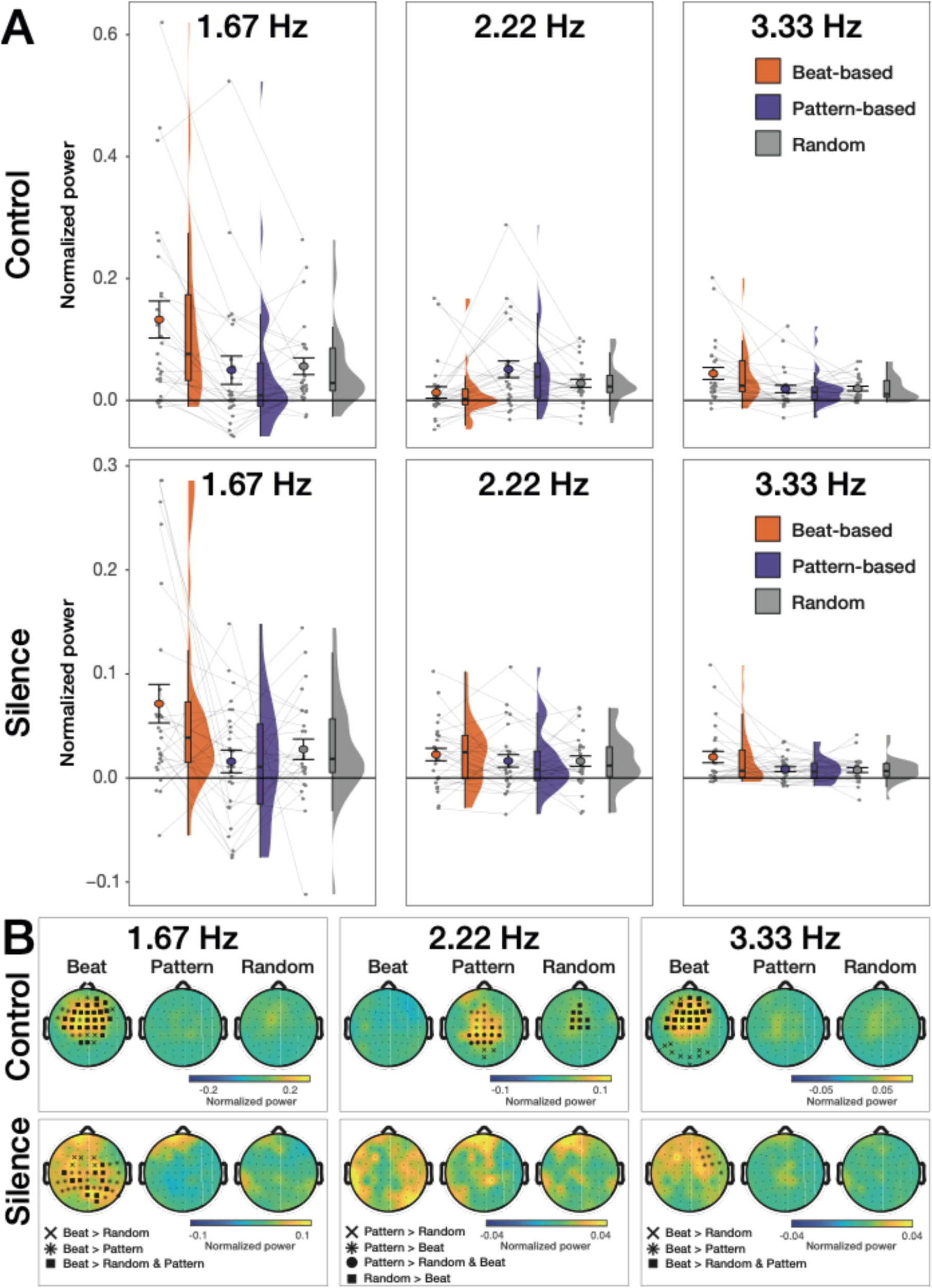
Individual participant data shows heterogeneity of the effects of temporal expectations on spectral power. All plots depict the spectral power at the frequency of interest, averaged over all electrodes, and normalized to account for the 1/f distribution (see Materials and Methods). A) Single participant data, with all data points in grey. Boxplots show the median, with the lower and upper hinges corresponding to the first and third quartiles (25^th^ and 75^th^ percentiles), and the whiskers corresponding to values no further than 1.5 times the inter-quartile range from the hinges. Error bars depict 2 standard errors around the mean. B) Scalp topographies for all conditions. Electrodes contributing to significant clusters in the cluster-based tests are highlighted on the plots for the conditions from each comparison in which power was largest.

Importantly, during the silence (Figure 6, right, and Figure 7, bottom), no significant clusters were found at 2.22 Hz, but at the beat frequency (1.67 Hz), power was significantly larger for the beat-based than the pattern-based (*p* = 0.01) and random (*p* = 0.012) conditions. In addition, at 3.33 Hz, power for the beat-based condition was larger than for the pattern-based condition (*p* = 0.038), with a trend when comparing the beat-based with the random condition (*p* = 0.077). Thus, while the pattern-based and random conditions showed tracking of the sound during stimulation (which is sometimes considered entrainment “in the broad sense” (Obleser & Kayser, 2019)), in the silence, entrainment was only present for the beat-based condition. This finding fits our behavioral observations that differentiate qualitatively between the effects of beat-based expectations and pattern-based expectations in the silence. To further substantiate the absence of persistent entrainment at the pattern frequency in the silence, we performed a Bayesian T-test comparing the normalized power at 2.22 Hz between the pattern-based and random conditions. We found moderate evidence in favor of the null hypothesis (no difference between conditions) (BF_01_ = 4.5). The results did not change as a function of the prior used (with a more traditional prior of r = 1, BF_01_ = 6.17).

Note that like for the behavioral results indicative of entrainment, there was large heterogeneity between participants (see Figure 7). While the power differences were significant in the overall cluster-based analyses, out of 27 participants, only 16 showed on average (i.e. over all electrodes) numerically larger power in the beat-based condition at the beat frequency when compared to both the random and pattern-based condition.

### Multiscale entropy as a non-stationary measure of temporal expectations

Figure 8 shows sample entropy for each condition separately, as well as the electrodes contributing to significant clusters in the analysis. Given that MSE indexes signal irregularity, we would expect entropy to be higher for the random and pattern-based conditions than for the beat-based condition in the silence. A cluster-based test on all electrodes and timescales showed that both in the control window, and in the silence window, entropy was higher for the pattern-based than for the random condition (control: *p* = 0.03; silence: *p* = 0.021). Note that in the silence window, all but the two highest timescales were included in the cluster. In the control window, the cluster spanned all timescales from 35 till 406 ms (see Materials and Methods for an explanation of the timescales). These results suggest that the signal was more irregular in the pattern-based than the random condition, over a broad range of timescales. Neither the difference in entropy between the beat-based and pattern-based condition, nor between the beat-based and random condition was significant, in either control or silence windows (all *p*s > 0.24).

**Figure 8.**
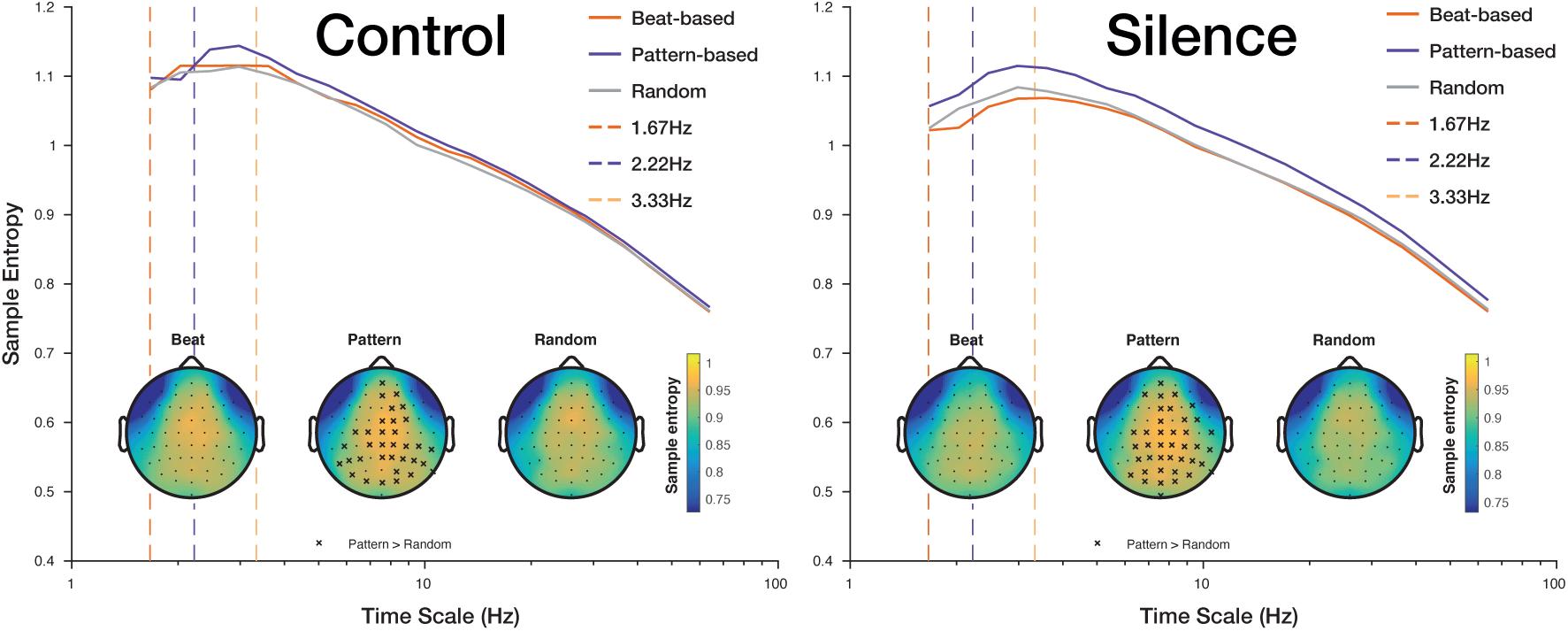
Entropy was higher for pattern-based than random sequences in both control and silence windows. Entropy in the Control window (left) and Silence window (right), averaged over frontocentral electrodes, and scalp distributions averaged over all timescales, depicting electrodes contributing to significant clusters.

Entropy has been related to various other EEG measures, such as spectral power and overall differences in signal variability (Kosciessa et al., 2020). To account for these, we ran several additional analyses. First, to check whether the differences in signal variability may have been caused by differences in low frequency activity, we repeated the MSE analysis on high-pass filtered data (Kloosterman et al., 2020), using a 5 Hz high-pass filter. This completely removed the effects (all *p*s > 0.43), suggesting that the differences between conditions were caused by low frequency activity in the signal (see Figure A2 in the Appendix).

Second, we checked whether differences in signal variability caused the differences in entropy, since entropy is calculated relative to the overall signal standard deviation (e.g., a pattern is considered a match at a lower threshold when overall signal variability is high). As can be seen in the Appendix (Figure A3), the similarity bounds used to compute entropy (derived from the time-domain signal standard deviation) differed between conditions, and this difference mirrored the differences in entropy, suggesting that at least some of the variance we observed was due to overall signal variability, and not necessarily signal irregularity.

### Multivariate decoding as a time-resolved method for studying entrainment

With multivariate decoding, we expected that training at expected times would yield above chance performance when testing at expected times, regardless of whether these time points were the same (e.g., when training at 600 ms, we expected to be able to accurately distinguish the beat-based from the random condition when testing at not just 600 ms, but also at 1200 and 1800 ms, as all these times were on the beat, or similarly expected). Figure 9A shows the temporal generalization matrices for each comparison, with significant clusters indicated by a black contour. In the silence window we found above chance decoding when decoding the beat-based against the random condition (*p* = 0.043), the pattern-based against the random condition (*p* < 0.001), and the beat-based and pattern-based conditions against each other (*p* = 0.009), indicating that based on the EEG signal in the silence, we could classify which type of rhythm participants had heard just before. However, looking at the temporal generalization matrices, it becomes apparent that contrary to our expectations, this above-chance decoding was not due to recurrent activity for expected events. Only clusters on the diagonal were significant for each comparison. In addition, decoding for all three comparisons was best in the second halve of the silence window, which is where the probe tones were presented. This suggests that the decoding mainly picked up on task-related differences. While the probes were physically identical across conditions, participants may have had different strategies to perform the task, depending on the type of sequence, and this could have resulted in above-chance decoding related to the task and probes, even when only analyzing the silence window.

**Figure 9.**
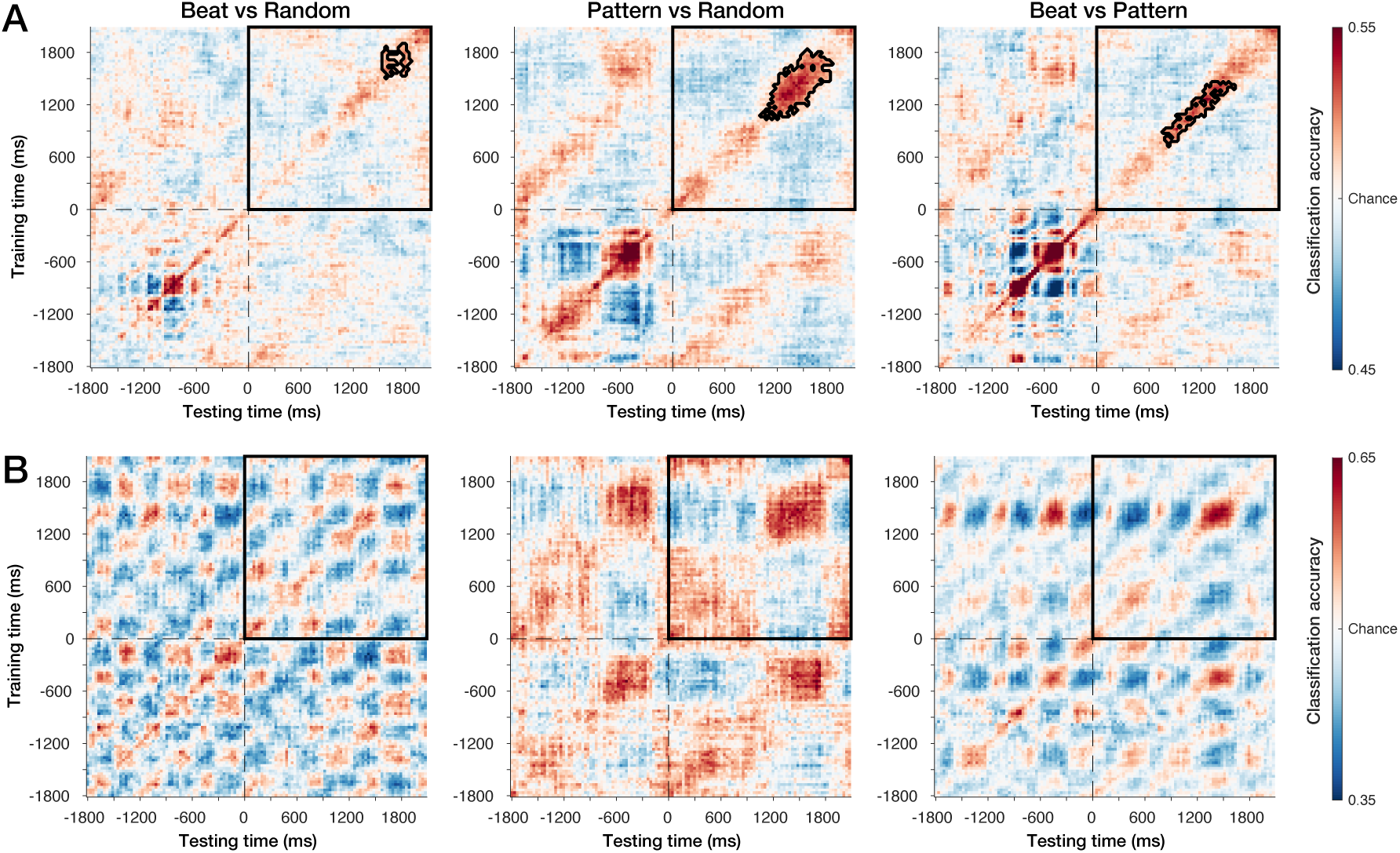
No recurrent activation in group average decoding, example of recurrency in single participant. A) Temporal generalization matrices for each comparison for the group analysis. The Y-axis shows the training time points, and the X-axis testing time points. The color scale indicates the classification accuracy, with 0.5 being chance level (all analyses are based on decoding two conditions against each other). Significance of classification accuracy was assessed by using cluster-based permutation tests on the silence window (300-2100 ms after the last note onset). The black contour indicates significant clusters. Note that the upper right quadrant, highlighted by the black box, is the entire silence window. B) Temporal generalization matrix for a participant showing recurrent activity following the expected beats. When decoding the beat-based vs. the other two conditions (left- and rightmost plots), peaks in accuracy follow a clear oscillatory pattern, with a phase consistent with the beat-based sequence (600 ms between beats and decoding peaks).

Of note, the decoding results varied considerably between participants. In one participant in particular (Figure 9B), we observed a pattern in the temporal generalization matrix that was consistent with the hypothesized result for an oscillatory process (King & Dehaene, 2014). However, even though some other participants also showed recurrent activity, the exact times of most accurate decoding, and the exact period of the recurrent process, differed widely between participants, obscuring these effects in the grand averages. These individual differences may be caused by individual preferences for a level of regularity in the beat-based stimuli (Drake et al., 2000), with some people attending mostly to the beat level (1.67 Hz), but others possibly attending more to subdivisions (3.33 Hz) or the level of the meter (0.83 Hz). Also, for different people, the optimal phase of delta oscillations (i.e., the phase that aligns with expected moments) may differ (Breska & Deouell, 2017; Henry & Obleser, 2012), possibly causing optimal decoding at different time points. To circumvent the large task-related effects apparent in the decoding results, and the possible individual differences in phase and metrical level attended to, we ran an additional exploratory decoding analysis looking at decoding within instead of between conditions. Here, we found above-chance decoding of beat positions against offbeat positions only following the beat-based condition. However, while decoding was above chance in the beat-based condition, it was in fact not better than in the other conditions. As such, these results provide only weak support for persistent effects of beat-based, but not pattern-based expectations. Detailed results of the within-condition decoding analysis can be found in the Appendix (Figure A4).

## Discussion

In the current study, we aimed to directly compare the neural mechanisms underlying temporal expectations based on regular beats and predictable patterns, by examining the development of climbing activity and the persistence of neural entrainment after cessation of rhythmic input. Instead of isochronous sequences, which can elicit both beat-based and memory-based expectations, we used varying non-isochronous patterns to separate beat-based and pattern-based expectations. Moreover, by assessing responses in silence, we sidestepped possible confounds associated with acoustic differences between conditions (Capilla et al., 2011; Haegens & Zion Golumbic, 2018; Novembre & Iannetti, 2018; Zoefel et al., 2018). First, behaviorally, we found that while the effects of beat-based expectations spanned two beat cycles, the effects of pattern-based expectations reflected primarily the first expected moment in time. Second, a negative ERP component at around 300 to 450 ms after the onset of the last sound was larger for pattern-based than beat-based sequences. Third, we observed more power at the beat frequency in the silence window when participants were previously listening to a beat-based sequence as compared to a pattern-based or random sequence. Taken together, these findings suggest that beat-based and pattern-based expectations may rely on different neural mechanisms, as previously found for cue-based expectations (Breska & Deouell, 2017), and not shared mechanisms, be it entrainment (Tichko & Large, 2019), or a general top-down mechanism (Rimmele et al., 2018). Tentatively, our results could be interpreted as entrainment underlying beat-based, but not pattern-based expectations, with pattern-based expectations relying on a different mechanism, such as climbing neuronal activity.

Behaviorally, for the beat-based condition, we observed an inverted U-curve as late as 1200 ms after the final tone of each sequence in both experiments, in line with entrainment models (Bauer et al., 2015; Jones et al., 2002) that assume oscillations are self-sustaining (Large, 2008; Large & Palmer, 2002). Expectations in the beat-based condition led to both higher ratings for expected and lower ratings for unexpected events compared to the random condition. Divesting resources from times at which an event is unlikely may be metabolically beneficial (van Atteveldt et al., 2015), and as such, has been suggested as a hallmark of entrainment and the associated “rhythmic” mode of processing (Schroeder & Lakatos, 2009b; Zoefel & Vanrullen, 2017). Indeed, the unexpectedness of events off the beat has been proposed to be a better indication of beat-based expectations than the expectedness of events on the beat (Bouwer et al., 2020; Breska & Deouell, 2017), in line with the current results, where the effects of beat-based expectations at unexpected time points exceeded those at expected time points.

Pattern-based expectations led to higher ratings at the first expected time point, and lower ratings at the first unexpected time point, showing that participants did learn the pattern. However, the behavioral effects of the predictable pattern only reflected the first expected moment, but not the subsequent structure. One speculative explanation for this is that listeners form expectations following temporal patterns one interval at a time (e.g., in the absence of an event, no next interval is predicted), and in a probabilistic way (Cannon, 2021; van der Weij et al., 2017). Possibly, the expectation for an event at 780 ms led to an inverted U-shape in responses, similar to the inverted U around a beat, but with a wider distribution, and only occurring once. This could explain the ratings being equal for probes at 780 and 885 ms, both within a time window where a tone could be expected, and ratings being lower for probes at 1200 and 1485 ms, which fell further from the expected time. An alternative explanation could be that the repetitiveness of the pattern lead participants to use a different heuristic during the pattern-based sequences to guide their ratings, considering only one interval at a time, while the varying rhythmic pattern of the beat-based condition induced a strategy whereby participants were more inclined to consider positions after the first expected tone.

In line with entrainment models of beat-based expectations (Haegens & Zion Golumbic, 2018; Henry & Herrmann, 2014; Large & Jones, 1999; Obleser & Kayser, 2019), we found that EEG power at the beat frequency (1.67 Hz) and its harmonic (3.33 Hz) in the silence window was larger following beat-based than pattern-based or random sequences. Such enhanced power was not found for a frequency inherent to the pattern-based sequence (2.22 Hz). Ongoing oscillations in silence, after sensory input has stopped, are regarded as strong evidence for entrainment (Breska & Deouell, 2017; Haegens & Zion Golumbic, 2018; Obleser & Kayser, 2019; van Bree et al., 2021; Zoefel et al., 2018). Our observation of enhanced power at the beat frequency during silence thus provides converging evidence for the notion that entrainment of low-frequency neural oscillations underlies beat-based perception, and our design, using non-isochronous rhythms, allows us to separate beat-based aspects from other structures present in natural rhythm (Bouwer et al., 2021), similar to other studies combining frequency tagging with non-isochronous rhythms (Lenc et al., 2021; Nozaradan et al., 2012; Tal et al., 2017).

Previously, no evidence for persistent entrainment at the frequency of isochronous rhythmic stimulation was found (Pesnot Lerousseau et al., 2021). Possibly, when presented with stimuli that do not require beat-based expectations to track the temporal structure, the brain may not engage in forming such expectations. Also, while in our study, the auditory sequences were task relevant, and the task itself was rhythm-related, in the study by Pesnot Lerousseau et al. (2021), participants listened to rhythms passively. Thus, the presence of persistent entrainment may also depend on task demands (Shalev et al., 2019). The influence of contextual factors on entrainment, like stimulus and task, are important topics for future research, as is the influence of movement. While we instructed participants not to move to the sequences, we cannot rule out that small, unintentional movements may have affected the EEG signal. More generally, movement may affect rhythm processing (Su & Pöppel, 2012), and thus entrainment.

The presence of persistent entrainment only in active listening conditions may suggest that beat-based expectations result from an explicit, top-down guided process, rather than some passive, automatic one (Bouwer, 2022). Technically, at the scalp, phase locking due to sustained oscillations and phase locking due to recurrent top-down signals may be indistinguishable, just as phase locking during rhythmic stimulation is often confounded with contributions from tone-evoked responses (Capilla et al., 2011; Novembre & Iannetti, 2018; Zoefel et al., 2018). Interestingly, in our study, the scalp topographies for power at 1.67 Hz differed between the silence and control windows, with power in the control window largest above a frontocentral region, and in the silence window above a parieto-central region. This may suggest that instead of resulting from a sustained automatic oscillation that would have a similar source during sound presentation and silence, persistent entrainment could originate from a different source, such as explicit predictions made by a motor network (Rimmele et al., 2018). Our study raises two questions in this regard, notably whether such a top-down signal would function differentially for beat-based and pattern-based expectations, as suggested by our results, and whether such a top-down signal would be related to CNV-like climbing neuronal activity.

The absence of power at 2.22 Hz following the pattern-based sequences suggests that entrainment does not underlie expectations based on learning a pattern, contrary to a recently proposed oscillator model that captures aspects of pattern-based expectations (Tichko & Large, 2019). In the time-domain, we found a negative deflection in the silence window following the pattern-based and random sequences, akin to a CNV. However, while previous studies showed a CNV peaking at an expected time (Breska & Deouell, 2017; Mento, 2017; Praamstra et al., 2006), here, the peak latency of the negative deflection in the signal was earlier, peaking around 400 ms for the pattern-based condition, while the first expected time point in the silence was at 780 ms. This may be explained by assuming that the CNV indexes temporal expectations in a probabilistic, and context-dependent way (Capizzi, Correa, & Sanabria, 2013; Damsma et al., 2021; Los & Heslenfeld, 2005; see also Hassall et al., 2022). The pattern used in the current study contained temporal intervals with durations between 150 and 780 ms. The ERP peaking at 400 ms may have indexed the average interval presented in the sequence (∼360 ms). This explanation is supported by the presence of a similar ERP deflection for the random condition, which while random in terms of transitional probabilities, was identical to the pattern-based condition in terms of the absolute intervals used. Of note, probabilistic models indexing statistical regularities in inter-onset intervals have indeed been used to explain aspects of temporal processing (Cannon, 2021; Elliott et al., 2014; van der Weij et al., 2017). In future work, linking such models directly to neural markers of pattern-based expectations may provide more insight in the mechanisms underlying pattern-based expectations and how they relate to the CNV.

Interestingly, the same ERP component was less present in the beat-based condition, despite an identical average duration of intervals. In previous research, a CNV was found peaking at expected times not only for cue-based, but also for beat-based expectations (Breska & Deouell, 2012, 2017; Breska & Ivry, 2020; Praamstra et al., 2006), though typically, with isochronous stimulation. Therefore, memory-based expectations, be it pattern-based or cue-based, could also have been formed in response to these sequences (Bouwer et al., 2016, 2020, 2021; Breska & Ivry, 2016; Keele et al., 1989), and may have contributed to the elicitation of a CNV. Here, using a beat-based sequence that did not allow for expectations based on simply learning transitional probabilities, we did not observe the same negative deflection as in the pattern-based sequences. This raises the possibility that a mechanism related to climbing neuronal activity (or CNV) may specifically support the formation of pattern-based and cue-based (Mento, 2013, 2017), temporal expectations, and that if possible given the input, the brain operates in a rhythmic mode of processing (Rimmele et al., 2018; Schroeder & Lakatos, 2009b).

It could be argued that the differences we observed in ERPs were caused by differences in the P3 response to the last sound of each sequence rather than the CNV, which would be apparent at a similar latency. The P3 is indeed susceptible to temporal expectations, with larger amplitude responses for temporally predictable targets (Lange, 2009; Mento, 2017; Schmidt-Kassow, Schubotz, & Kotz, 2009). However, these effects can be observed for beat-based and memory-based expectations in a similar direction (Breska & Deouell, 2017; Breska & Ivry, 2020; Mento, 2017; Schmidt-Kassow et al., 2009). Hence the P3, if anything, should have been larger for the pattern-based than random sequences, which is not the case. Thus, we feel it is unlikely that the ERP differences are caused by differences in the P3, and we tentatively suggest that here, the ERP results are more likely to be due to a CNV-like mechanism.

One alternative explanation for our results could be that participants used an interval-based strategy in the beat-based condition, predicting an event every 600 ms, and that the differences between conditions were due to easier learning in the beat-based than pattern-based condition, both because the length of the beat (600 ms) was shorter than the length of the pattern (1800 ms), and because the perceived beat occurred three times as often as the pattern. However, first, because of the varying surface structure in the beat-based condition, in absolute terms the beat interval occurred exactly as often as the pattern intervals. Second, the pattern was kept consistent throughout the experiment, for a total of 216 presentations, making it unlikely that participants did not learn the pattern. Indeed, a previous study showed that even untrained participants could reproduce non-metrical patterns to some extent after only one presentation (Cameron & Grahn, 2014). Third, in our previous study, using the same stimuli, we showed that the behavioral facilitation caused by introducing a regular beat was smaller than the facilitation caused by introducing a predictable pattern (Bouwer et al., 2020), regardless of differences in length and number of presentations. Finally, all participants except one were able to judge the first probe after the pattern-based sequences as not fitting the pattern, as apparent from lower ratings for this probe in the pattern-based than random condition. This is comparable to the results for probes at unexpected times in the beat-based condition. Hence, differences in ease of learning are unlikely to explain our results.

In addition, listening to strictly metric patterns, as the ones used here, is associated with activity in a circuit including the basal ganglia, while listening to non-metric patterns is associated with activity in a circuit including the cerebellum, making it unlikely that the same, interval-based mechanism would be used for both types of rhythms (Leow & Grahn, 2014). Also, predicting the timing of events in non-isochronous strictly metric sequences would require participants to learn not just the transitional probabilities of single intervals, but also to combine multiple intervals into groups that together last the length of a beat. It is currently unclear whether humans, when faced with rhythmic patterns, use such a hierarchical interval-based strategy. Indeed, future research could examine if beat-based expectations in general can be explained by such multilevel interval learning, akin to a recent model for beat-based perception (Cannon & Patel, 2021), as this could provide a general challenge to oscillator models of beat-based perception. In such a view, the difference between beat-based and pattern-based timing may be the importance of hierarchical structure in beat-based rhythms (Fitch, 2013), rather than the presence of oscillations.

In addition to behavioral, ERP, and frequency-domain analyses, we looked at multi scale entropy and multivariate pattern analysis as alternative ways to examine neural entrainment. At the group level, entropy was higher following the pattern-based than random sequences. This could be explained by assuming that the brain uses a vigilance mode to track the pattern-based regularities, which at the neural level, translates in more irregular patterns of activity. Attention and arousal, which are both associated with temporal expectations (Schroeder & Lakatos, 2009a), have indeed been linked to neural variability as well (Waschke et al., 2021). However, the entropy results at least to some extent were explained by overall differences in signal variability (e.g., the signal variance), so this hypothesis remains to be confirmed in future research. Considering the decoding results, the observed above-chance decoding seemed to primarily reflect general task-related activity. The best decoding was observed in the second half of the silence window, where probes could be presented. Also, in the group-average decoding results, above chance decoding was limited to training and testing on the same time points, and we did not observe recurrent activity. Yet, we did show a proof of concept for our approach in at least one participant, who showed a clear oscillatory pattern in decoding accuracy when decoding the beat-based against the other two conditions. This pattern of activity is in line with the strength or sharpness of the neural representation of tones varying over time as a function of temporal expectations (Auksztulewicz et al., 2018, 2019).

The strength of the decoding approach is that neural dynamics can be assessed in real time, but this may also be its weakness, particularly in the current study. Due to the absence of sensory input, individuals may differ in how consistently they persist in their beat percept. Also, people may attend to different levels of regularity in beat-based perception (Drake et al., 2000), and the optimal phase of delta oscillations (e.g., the phase that aligns with expected moments) may differ across individuals (Breska & Deouell, 2017; Henry & Obleser, 2012; Sun et al., 2021). One option to overcome the weaknesses of the decoding approach could be to transform the EEG signal to allow frequency and phase drift. Recent technical implementations of time warping algorithms may be fruitful in this regard (Chemin et al., 2018; van Bree et al., 2022).

The more traditional frequency tagging approach allows for an assessment of activity across multiple frequencies, as present in rhythm in a metrical structure. Phase-based analyses may not be suitable to answer the questions we ask here, due to differences in preferred phase between individuals, as well as the fact that phase may not differentiate between beat-based and memory-based expectations (Breska & Deouell, 2017). However, a frequency tagging approach is not sensitive to phase differences between individuals. This may explain why the results for the frequency-based analysis were more robust than those obtained with decoding. A weakness of the frequency tagging approach may be that the Fourier transform assumes stationarity in the oscillating signal, while entrainment models propose a dampening factor to account for decreasing oscillatory power over time (Doelling & Assaneo, 2021; Large et al., 2015). To assess power at specific frequencies in a time-resolved way, wavelet convolution is often used as an alternative. But, in the current study, differentiating between the specific frequencies of the beat and the pattern would require wavelet parameters that would result in a temporal resolution too low to disentangle activity during and before the silence (i.e., many wavelet cycles would be needed). Recently, a promising alternative to assess oscillatory activity in the time domain was proposed in the form of cycle-by-cycle analysis (Cole & Voytek, 2019). However, this approach requires filtering the signal at the specific frequency of interest, again posing problems for disentangling low frequency oscillations due to the beat, the pattern, and ongoing ERPs. Assessing the time course of low frequency oscillations thus remains a challenge for future research.

In our current study, we observed large heterogeneity between individuals. While it is often assumed that most people automatically form beat-based expectations (Honing, 2012), recent evidence showed phase locking to speech in only about half of the population (Assaneo et al., 2019). Indeed, in our study, only about two-thirds of the participants behaviorally showed evidence for beat-based expectations in the second half of the silence window and we only observed enhanced power at the beat frequency following beat-based sequences in about half of the participants. The behavioral effects of beat-based and pattern-based expectations were also associated with musical training, consistent with previous research using beat-based (Bouwer et al., 2016, 2018; Cameron & Grahn, 2014; Matthews et al., 2016; Vuust et al., 2005) and pattern-based (Cameron & Grahn, 2014) rhythms. Some previous studies have failed to show differences between musicians and non-musicians, however (Bouwer et al., 2014; Geiser et al., 2009; Grahn & Brett, 2007), possibly due to differences in task design (Bouwer et al., 2018). The current study used an explicit rating task, for which performance may be particularly improved by musical training, as musically trained participants may have additional strategies to perform the task. The use of implicit timing tasks may be a better probe of innate differences in timing abilities, which need not necessarily be related to musical training (Law & Zentner, 2012), and implicit tasks may also be less susceptible to task-related effects as we observed in the decoding analysis, which may stem from individual strategies in performing the explicit task. Undoubtedly, given the heterogeneity we and others (Assaneo et al., 2019; Bauer et al., 2015; Sun et al., 2021) have observed in tasks probing temporal expectations, understanding individual differences is an important direction for future research, with significant implications for applications of musical rhythm, such as in motor rehabilitation (Dalla Bella et al., 2018).

## Conclusion

In summary, we have shown that beat-based and pattern-based expectations can be differentiated in terms of their behavioral and neurophysiological effects once sensory input has ceased. These findings provide novel evidence for the notion that different mechanisms implement temporal expectations based on periodic and aperiodic input streams, with, tentatively, the former based on entrainment of low frequency neural oscillations, and the latter on climbing neural activity indexing a memorized interval.

## Appendix

### Multi Scale Entropy computation

To compute entropy, epochs were concatenated. Sample entropy was then calculated by taking the following steps:

1. In each time series, a template is selected, consisting of *m* samples. The calculation of entropy is an iterative process, using each sample in the time series as a starting point for the template once.
2. Throughout the time series, the algorithm searches for patterns that match the template pattern. A section of samples is considered a match if it resembles the template pattern enough to fall within a set boundary, which is defined as *r* x SD (the similarity bound).
3. The number of pattern matches is counted.
4. Subsequently, the same procedure is followed for patterns of *m* + 1 samples long.
5. For the total counts of pattern matches throughout the time series, sample entropy is then calculated as the logarithm of the ratio between pattern matches of length *m* and pattern matches of length *m* + 1.

Thus, sample entropy reflects the proportion of patterns in the time series that stays similar when an extra sample is added to the pattern. Here, we used *m* = 2 and *r* = 0.5, as was done previously for EEG data (Kloosterman et al., 2020).

Sample entropy is then repeated for multiple timescales, to account for contributions of both low and high frequency neural activity. The time series is coarsened step by step, by taking the average of a group of adjacent samples with step-wise increasing group size. This means that long, or coarse, timescales are equivalent to low frequency activity. For example, at a sampling rate of 256 Hz, a timescale of 4 (e.g., averaged over four adjacent samples, or 15.6 ms) is roughly the equivalent of looking at activity at 64 Hz, while a timescale of 153 at that sample rate corresponds to averaging over 598 ms, or the equivalent of activity at roughly 1.67 Hz. Here, we used twenty timescales ranging from 4 till 153, with 153 being the maximum timescale given the length of an epoch of 1800 ms (e.g., one epoch equals 460 samples, but since entropy is calculated on patterns of 2 and 3 samples, the maximum coarsening to retain the possibility of having a 3-sample pattern is by averaging over 153 samples). Similarity bounds were recomputed for each time scale (Kloosterman et al., 2020; Kosciessa et al., 2020). To control for the contribution of delta, we repeated the analysis on high-pass filtered data (5 Hz).

### Decoding within condition

The initial decoding analysis yielded large effects that may be task-related, and individual differences in phase and metrical level attended to may have additionally hampered finding condition differences at the group level. Therefore, we ran an additional decoding analysis in which we decoded expected and unexpected positions against each other within each condition. For this analysis, we only included frontocentral electrodes (C1, C2, C3, C4, Cz, FC1, FC2, FC3, FC4, FCz, F1, F2, F3, F4, Fz), which were the same electrodes that showed the largest P1 responses to the initial sounds of the sequences, indicative of representing the auditory cortex. Here, we defined time points in the silence as expected or unexpected based on the rhythmic sequences. To equate the choice of expected and unexpected time points as much as possible in terms of number of time points included and distance between time points, we used times that were also used in the probe tone experiment: 600, 780, 885, and 1200 ms. For beat-based sequences, time points 600 and 1200 were expected (on the beat), while time points 780 and 885 were unexpected (offbeat). The same time points take on a different meaning if preceded by the pattern-based sequences. Then, 780 and 1200 ms are considered expected (predictable based on the pattern), while both 600 and 885 ms are unexpected (unpredictable based on the pattern). For all conditions, we examined whether we could classify above chance whether a time window of 100 ms centered on the time point of interest was an expected or unexpected moment in time. As for the first decoding analysis, the data were resampled to improve signal to noise, here to 128 Hz to retain enough data points in the 100 ms window for analysis. After the decoding, we extracted the average classification accuracy for the 100 ms time window to test significance for each condition separately against chance level, using t-tests against 0.5, and between conditions in the silence, using a repeated measures ANOVA, with condition as an independent factor, a random intercept for participant, and decoding accuracy as the dependent variable.

**Figure A1.**
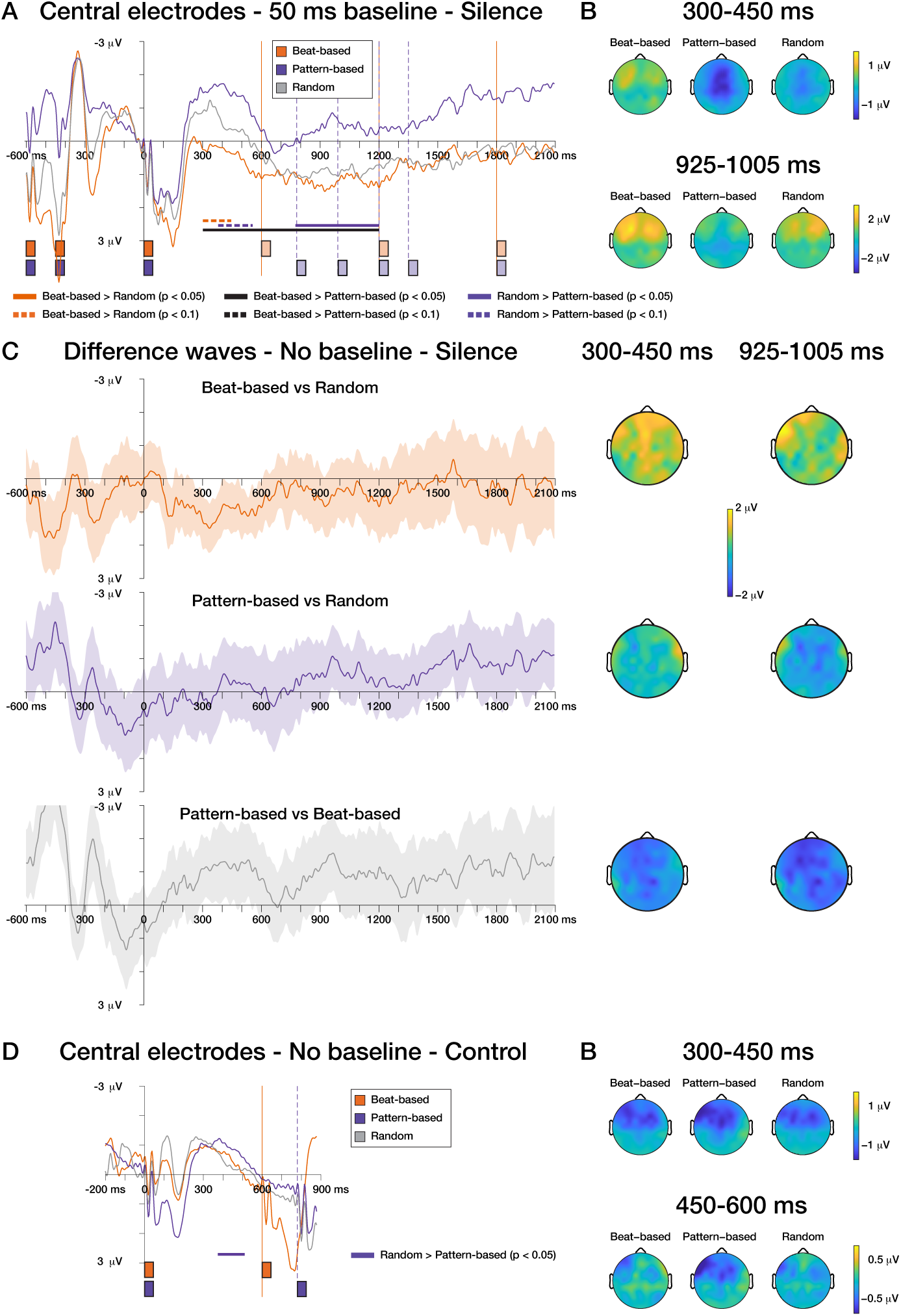
ERPs with traditional 50 ms baseline and difference waves. A) With the more traditional pre-cue baseline, there was a trend for the random condition to elicit a more negative deflection than the beat-based condition (*p* = 0.058, 300 – 446 ms) and for the pattern-based condition to elicit a more negative deflection than the random condition (early window: *p* = 0.083, 380 – 551 ms; later window: *p* = 0.022, 774 – 1200 ms). Note that a significant cluster spanning the entire analysis window of 300 – 1200 ms was found when comparing between the pattern-based and beat-based condition with baseline correction (*p* = 0.011), indicative of a possible overall shift due to the baseline correction. However, the overall picture stays the same as when not using a baseline, a negative deflection can be seen around 300-450 ms after the last sound onset for the pattern-based and random, but not the beat-based condition. B) Scalp distributions for windows with a significant cluster (see also Figure 5). C) Difference waves depicting condition differences and the 95% confidence interval around the mean difference waveforms. The scalp distributions similarly depict condition differences. D) Evoked potential showing the ERPs during sound presentation. Time 0 here is the presentation of a sound, with the next sound being presented at 600 (beat-based condition) or 780 ms (pattern-based and random condition). As in the silence window, a negative deflection can be observed, which was larger for the pattern-based than random condition (*p* = 0.039), and peaks around 350 ms after the onset of the previous sound. The difference between the pattern-based and beat-based condition here did not reach significance (*p* = 0.1), likely because of the noisy baseline.

**Figure A2.**
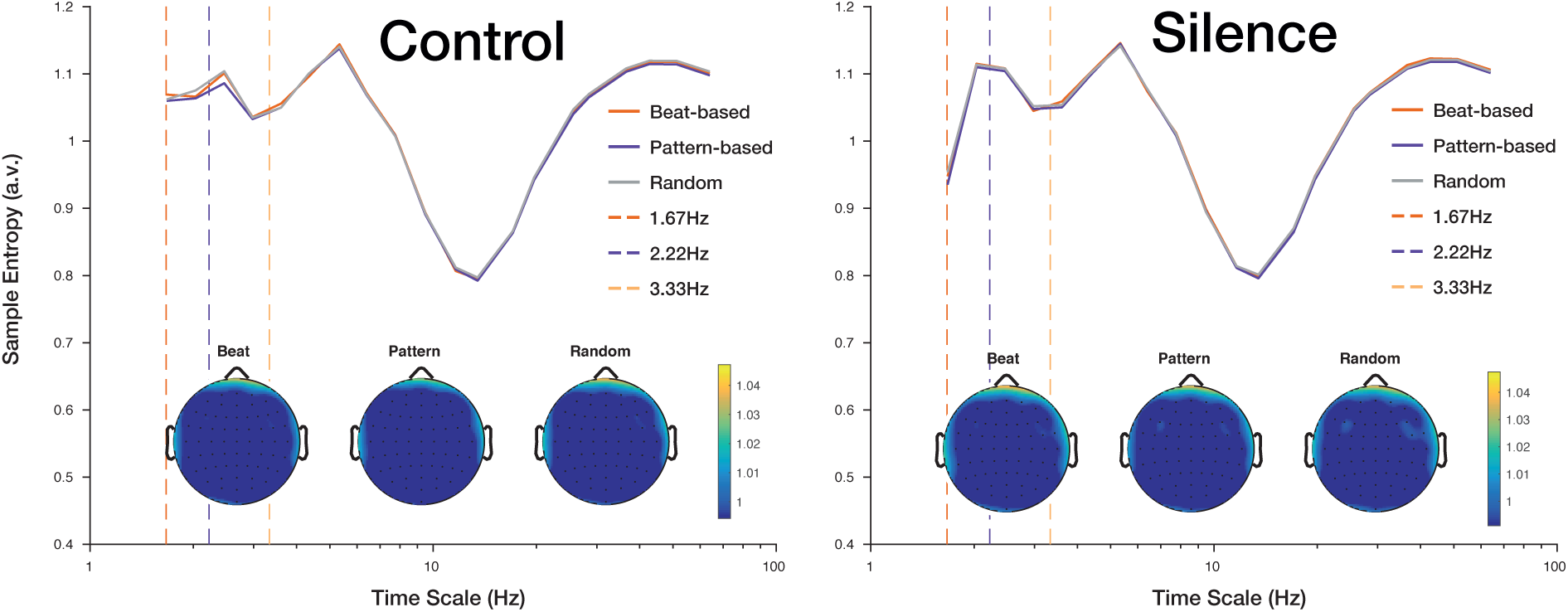
MMSE after highpass filtering at 5 Hz. Here, all differences between conditions are eliminated by the highpass filtering.

**Figure A3.**
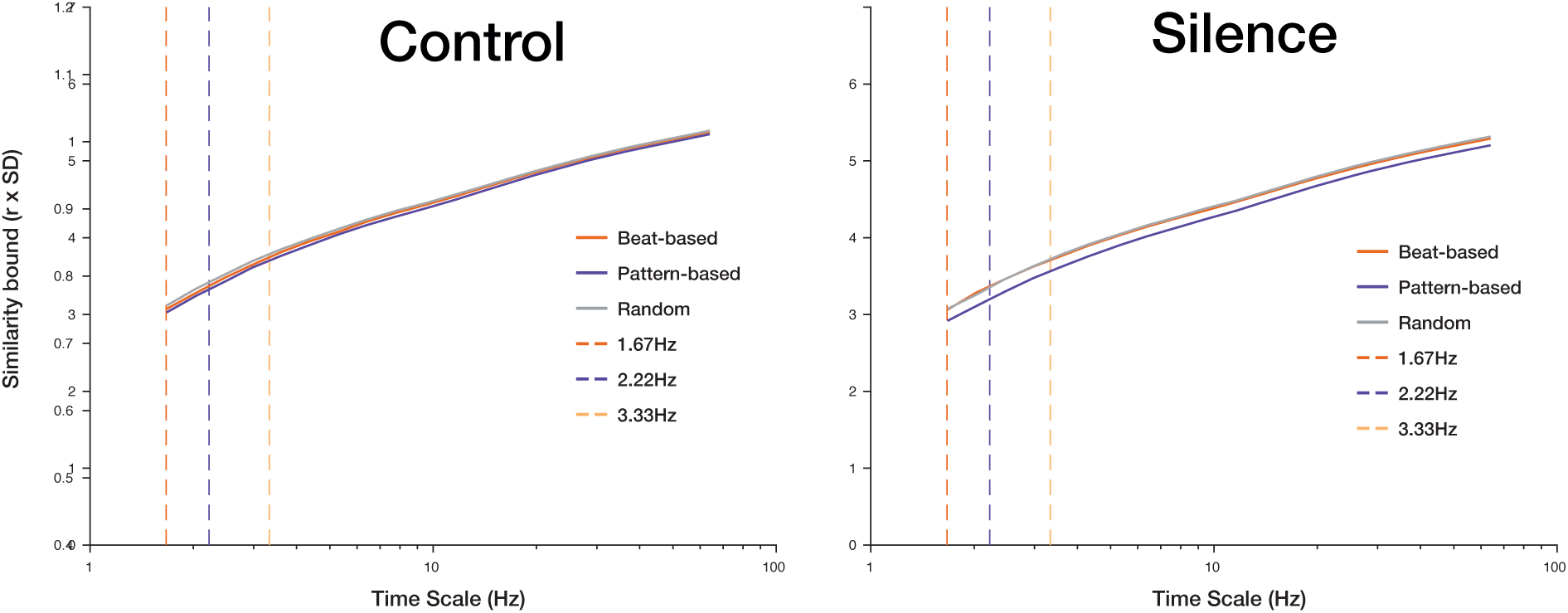
Similarity bounds used to calculate MMSE values mirror the condition differences. This finding suggests that condition differences found in the entropy measure, with higher entropy for the pattern-based than random condition in the silence window, may be due to overall variability differences, with lower variability for the pattern-based than random condition, rather than differences in entropy of the signal.

**Figure A4.**
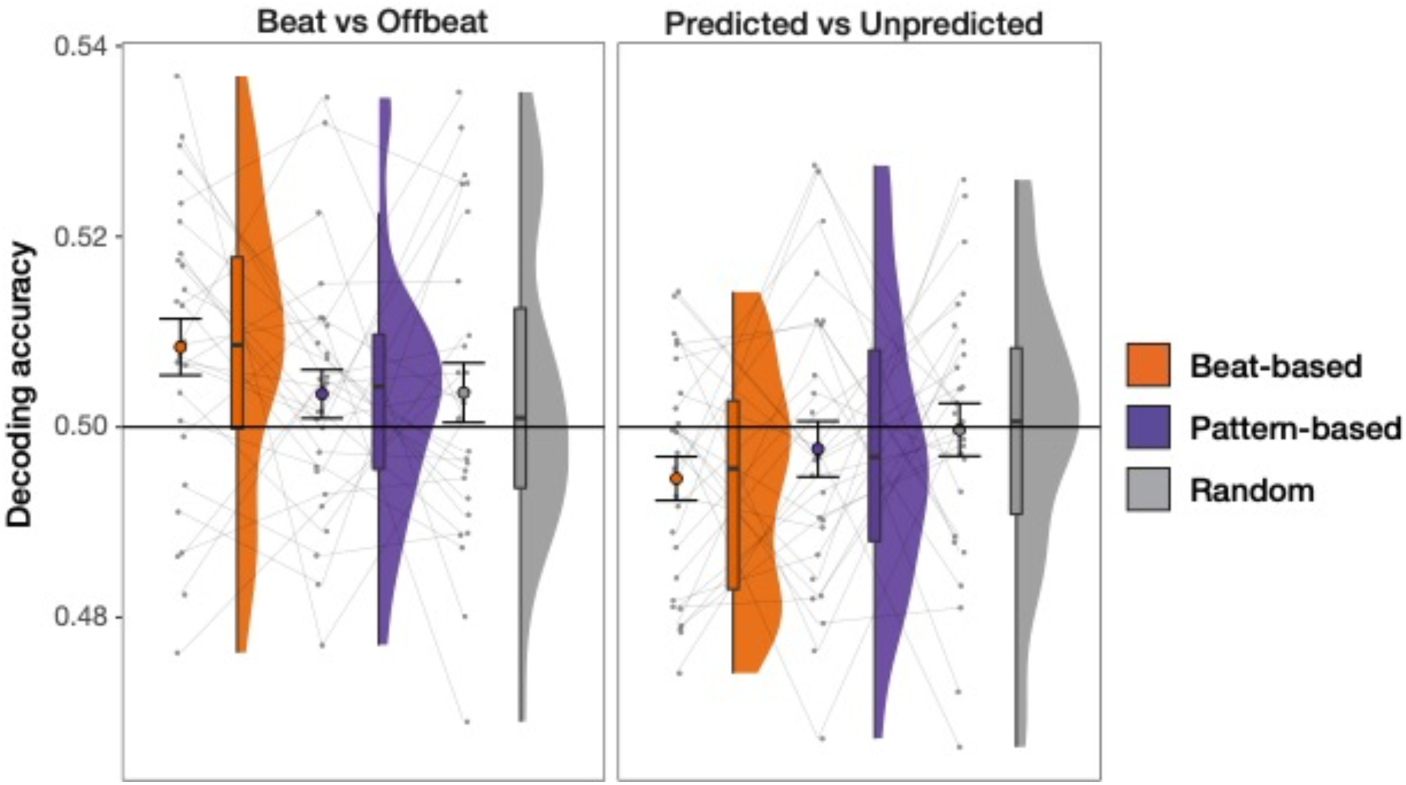
Above chance decoding for beat vs offbeat time points only after listening to beat-based sequences. For the within-condition decoding analysis, we used a classification algorithm to differentiate between times that were expected or unexpected based on the beat (on the beat: 600 and 1200 ms after the onset of the last tone; offbeat: 780 and 885 ms after the onset of the last tone) or pattern (predicted: 780 and 1200 ms after the onset of the last tone; unpredicted: 600 and 880 ms after the onset of the last tone). The graphs depict the average classification accuracy in a 100 ms time window centered on these expected and unexpected time points in the silence window. Here, the beat-offbeat comparison should yield better decoding if preceded by the beat-based sequences, while the predicted-unpredicted comparison should be decoded better if preceded by the pattern-based sequences. Only decoding of beat vs offbeat positions after the beat-based sequences yielded above-chance decoding (*t*_26_ = 2.84, *p* = 0.009). None of the other comparisons lead to decoding above chance (all *p*s > 0.18), but decoding of predicted vs unpredicted positions in the beat-based condition did yield below-chance decoding (*p* = 0.026). However, a subsequent ANOVA comparing decoding of beat against offbeat positions in all three conditions showed that decoding was not significantly better for the beat-based condition than the other conditions (effect of condition in the silence: χ^2^ = 2.1, *p =* 0.35). Like for the other analyses, there were larger differences between participants, and decoding, while above chance, did not exceed 0.55 accuracy. As for the other EEG measures, the difference in decoding accuracy between the beat-based and random condition, and the patten-based and random condition did not depend on musical training (both *p*s > 0.21).

**Table A1.** Simple effects per position from the full models, comparing three conditions.

|  |  | Experiment 1 |  |  |  | Experiment 2 |  |  |  |
| --- | --- | --- | --- | --- | --- | --- | --- | --- | --- |
| Position | Contrast | Estimate | SE | Z ratio | *p* | Estimate | SE | Z ratio | *p* |
| 600 ms | Random – Pattern | 2.4 | 0.09 | 26.2 | <.0001 |  |  |  |  |
|  | Random – Beat | -1.2 | 0.09 | 13.3 | <.0001 |  |  |  |  |
|  | Pattern – Beat | -3.5 | 0.09 | -38.3 | <.0001 |  |  |  |  |
| 780 ms | Random – Pattern | -0.3 | 0.09 | -2.9 | 0.058 |  |  |  |  |
|  | Random – Beat | 1.6 | 0.09 | 18.3 | <.0001 |  |  |  |  |
|  | Pattern – Beat | 1.8 | 0.09 | 20.7 | <.0001 |  |  |  |  |
| 885 ms | Random – Pattern | -0.1 | 0.09 | -1.6 | 1 |  |  |  |  |
|  | Random – Beat | 1.5 | 0.09 | 17.0 | <.0001 |  |  |  |  |
|  | Pattern – Beat | 1.6 | 0.09 | 18.4 | <.0001 |  |  |  |  |
| 990 ms | Random – Pattern | 0.1 | 0.09 | 1.1 | 1 | 0.6 | 0.1 | 4.8 | <.0001 |
|  | Random – Beat | 1.4 | 0.09 | 16.2 | <.0001 | 1.4 | 0.1 | 11.4 | <.0001 |
|  | Pattern – Beat | 1.3 | 0.09 | 15.0 | <.0001 | 0.8 | 0.1 | 6.5 | <.0001 |
| 1200 ms | Random – Pattern | 0.6 | 0.08 | 7.5 | <.0001 | 0.2 | 0.1 | 1.6 | 0.92 |
|  | Random – Beat | -0.3 | 0.09 | -3.7 | 0.004 | 0.1 | 0.1 | 0.9 | 1 |
|  | Pattern – Beat | -1.0 | 0.09 | -10.9 | <.0001 | -0.1 | 0.1 | -0.7 | 1 |
| 1485 ms | Random – Pattern | 0.5 | 0.09 | 5.2 | <.0001 | -0.5 | 0.1 | -4.4 | <.0001 |
|  | Random – Beat | 0.6 | 0.09 | 6.5 | <.0001 | 0.4 | 0.1 | 3.2 | 0.011 |
|  | Pattern – Beat | 0.1 | 0.09 | 1.3 | 1 | 0.9 | 0.1 | 7.5 | <.0001 |

*Note.* All *p*-values have been Bonferroni corrected for 18 (Experiment 1) or 9 (Experiment 2)
comparisons.

**Table A2.** Simple effects per condition from the baseline corrected model, using the random condition as baseline, for all positions.

|  |  | Experiment 1 |  |  |  | Experiment 2 |  |  |  |
| --- | --- | --- | --- | --- | --- | --- | --- | --- | --- |
| Condition | Contrast | Estimate | SE | Z ratio | *p* | Estimate | SE | Z ratio | *p* |
| Beat-based | 600 ms - 780 ms | 2.3 | 0.08 | 28.7 | <.0001 |  |  |  |  |
|  | 600 ms - 885 ms | 2.2 | 0.08 | 27.8 | <.0001 |  |  |  |  |
|  | 600 ms - 990 ms | 2.1 | 0.08 | 27.0 | <.0001 |  |  |  |  |
|  | 600 ms - 1200 ms | 0.6 | 0.08 | 6.9 | <.0001 |  |  |  |  |
|  | 600 ms - 1485 ms | 1.5 | 0.08 | 18.0 | <.0001 |  |  |  |  |
|  | 780 ms - 885 ms | -0.1 | 0.08 | -1.3 | 1 |  |  |  |  |
|  | 780 ms - 990 ms | -0.2 | 0.08 | -1.9 | 1 |  |  |  |  |
|  | 780 ms - 1200 ms | -1.8 | 0.08 | -21.3 | <.0001 |  |  |  |  |
|  | 780 ms - 1485 ms | -0.9 | 0.08 | -10.5 | <.0001 |  |  |  |  |
|  | 885 ms - 990 ms | -0.1 | 0.08 | -0.7 | 1 |  |  |  |  |
|  | 885 ms - 1200 ms | -1.7 | 0.08 | -20.3 | <.0001 |  |  |  |  |
|  | 885 ms - 1485 ms | -0.8 | 0.08 | -9.4 | <.0001 |  |  |  |  |
|  | 990 ms - 1200 ms | -1.6 | 0.08 | -19.5 | <.0001 | -1.1 | 0.1 | -9.6 | <.0001 |
|  | 990 ms - 1485 ms | -0.7 | 0.08 | -8.7 | <.0001 | -0.9 | 0.1 | -7.9 | <.0001 |
|  | 1200 ms - 1485 ms | 0.9 | 0.08 | 10.9 | <.0001 | 0.2 | 0.1 | 1.8 | 0.46 |
| Pattern-based | 600 ms - 780 ms | -2.3 | 0.09 | -27.0 | <.0001 |  |  |  |  |
|  | 600 ms - 885 ms | -2.2 | 0.08 | -26.4 | <.0001 |  |  |  |  |
|  | 600 ms - 990 ms | -2.1 | 0.08 | -24.8 | <.0001 |  |  |  |  |
|  | 600 ms - 1200 ms | -1.7 | 0.08 | -20.2 | <.0001 |  |  |  |  |
|  | 600 ms - 1485 ms | -1.7 | 0.09 | -20.3 | <.0001 |  |  |  |  |
|  | 780 ms - 885 ms | 0.1 | 0.08 | 1.0 | 1 |  |  |  |  |
|  | 780 ms - 990 ms | 0.2 | 0.08 | 3.0 | 0.080 |  |  |  |  |
|  | 780 ms - 1200 ms | 0.6 | 0.08 | 7.6 | <.0001 |  |  |  |  |
|  | 780 ms - 1485 ms | 0.6 | 0.08 | 6.9 | <.0001 |  |  |  |  |
|  | 885 ms - 990 ms | 0.2 | 0.08 | 2.0 | 1 |  |  |  |  |
|  | 885 ms - 1200 ms | 0.5 | 0.08 | 6.7 | <.0001 |  |  |  |  |
|  | 885 ms - 1485 ms | 0.5 | 0.08 | 6.0 | <.0001 |  |  |  |  |
|  | 990 ms - 1200 ms | 0.4 | 0.08 | 4.7 | <.0001 | -0.3 | 0.1 | -2.9 | 0.022 |
|  | 990 ms - 1485 ms | 0.3 | 0.08 | 4.1 | 0.001 | -0.9 | 0.1 | -8.3 | <.0001 |
|  | 1200 ms - 1485 ms | 0.0 | 0.08 | -0.5 | 1.00 | -0.6 | 0.1 | -5.4 | <.0001 |

*Note.* All *p*-values have been Bonferroni corrected for 30 (Experiment 1) or 6 (Experiment 2) comparisons.

## Data availability statement

All datafiles, and code used for data acquisition, data analysis, and figure creation, are available through https://osf.io/uwny8/.

## Author contribution

**Fleur L. Bouwer:** Conceptualization, Methodology, Formal analysis, Investigation, Writing – Original Draft, Writing – Review & Editing, Visualization

**Johannes J. Fahrenfort**: Software, Formal analysis, Writing – Review & Editing

**Samantha K. Millard**: Methodology, Formal analysis, Investigation, Writing – Review & Editing

**Niels A. Kloosterman**: Software, Formal analysis, Writing – Review & Editing

**Heleen A. Slagter:** Conceptualization, Methodology, Resources, Writing – Review & Editing, Supervision, Funding acquisition

## Acknowledgements

We would like to thank Babak Chawoush and Charlotte van Roijen for their assistance with the data collection, and fruitful discussions during the project.

